# An additional site mutation in MITA/STING gain-of-function mutants abolishes the autoimmune SAVI phenotypes and directs a therapeutic strategy

**DOI:** 10.64898/2026.01.03.697515

**Authors:** Fang-Xu Li, Sheng Liu, Zhi-Dong Zhang, Xin Shuai, Bi-Kun Xiao, Jia Yang, Defen Lu, Dandan Lin, Guijun Shang, Bo Zhong

## Abstract

STING-associated vasculopathy with onset in infancy (SAVI) is an autoimmune disease caused by gain-of-function mutations (GOFs) of MITA/STING and the most frequent GOFs for SAVI are MITA^N154S^ and MITA^V155M^. However, how MITA GOFs are spontaneously activated remains incompletely understood. Here, we show that the activity of MITA hinge-region GOFs is compromised by an additional mutation at Lys150 and that the SAVI phenotypes of MITA^N153S/WT^ mice are completely abolished in the MITA^K150N/N153S^ (MITA^NS/NS^) mice. Mechanistically, MITA GOFs constitutively associate with iRhom2 for the spontaneous ER-to-Golgi translocation, which is substantially inhibited by the introduction of a mutation at Lys150. Interestingly, cGAMP binds to MITA^NS^, triggers the ER-to-Golgi translocation of MITA^NS^ as well as the MITA^NS^-iRhom2 interaction, and induces the expression of downstream genes in *Mita*^NS/NS^ cells similarly as in *Mita*^+/+^ cells. Consistently, structural studies demonstrate an inactive open conformation of apo-MITA^NS^ characterized by connector region crossover and a curved filament of cGAMP-bound MITA^NS^ characterized by parallel connector regions, similar to those observed in wild-type MITA. Furthermore, we design a SAVI-inhibitory peptide (SIP) that selectively inhibits the interaction between MITA^N153S^ and iRhom2 and the activity of MITA GOFs and thereby abolishes the SAVI phenotypes of the MITA^N153S/WT^→WT chimeric mice. These findings reveal a previously uncharacterized mechanism for the spontaneous activation of MITA GOFs and highlight a potential therapeutic intervention for SAVI.

## 1. Introduction

Mediator of IRF3 activation (MITA, also known as STING, TMEM173, ERIS, and MPYS) is an endoplasmic reticulum (ER) cyclic di-nucleotide sensor that mediates cytoplasmic double-stranded DNA (dsDNA)-triggered signaling (Ishikawa and Barber, 2008; Ishikawa et al., 2009; Jin et al., 2008; Sun et al., 2009; Zhong et al., 2008). MITA/STING consists of 4 transmembrane (TM) domains in the N terminus, an intermediate ligand-binding domain (LBD), and a C-terminal tail (CTT) domain (Hussain et al., 2022). Cryo-electron microscopy (cryo-EM) analyses of human MITA reveal a hinge region (residues 141-156) consisting of a connector helix (residues 141-149) and a connector loop (residues 150-156) between the TM4 and the first helix in the LBD (LBDα1) (Shang et al., 2019). Two copies of the hinge region form a right-handed crossover that makes two MITA molecules as a dimer (Shang et al., 2019). The MITA dimers exhibit an autoinhibitory structure through head-to-head and side-by-side packing at the ER under homeostatic conditions (Liu et al., 2023). Upon the binding of cyclic GMP-AMP (cGAMP) that is generated by the cGAMP synthase (cGAS) after sensing of dsDNA (Sun et al., 2013; Wu et al., 2013), MITA dimers undergo conformational changes by inducing a 180° rotation of the LBD relative to the TM domain and thereby expose the CTT domain (Gao et al., 2013; Zhang et al., 2013, 2019). In addition, the TM domains and the LBDs of one cGAMP-bound MITA dimer interact with those of another neighboring dimer, respectively, which results in the formation of a curved monolayer filament of MITA (Ergun et al., 2019; Liu et al., 2023). Subsequently, the cGAMP-bound curved MITA filament deforms the ER membrane to support its ER exit and transportation to the Golgi apparatus where MITA binds to Golgi apparatus-synthesized sulfated glycosaminoglycans for further polymerization (Fang et al., 2021). The polymerized MITA filament exposes CTTs to recruit TANK-binding kinase 1 (TBK1) to trans-phosphorylate Ser366 of the CTT domain for further recruitment and phosphorylation of interferon regulatory factor 3 (IRF3), thereby inducing the expression of type I interferons (IFNs) (Liu et al., 2015; Tanaka and Chen, 2012).

Aberrant activation of MITA results in various autoimmune diseases (Decout et al., 2021; Miner and Fitzgerald, 2023; Zhong and Shu, 2022), one of which, termed STING(MITA)-associated vasculopathy with onset in infancy (SAVI), is caused by the gain-of-function mutations (GOFs) of MITA (Jeremiah et al., 2014; Liu et al., 2014). Patients with SAVI typically exhibit interstitial lung disease, skin lesions, vasculitis, recurrent fever, and systemic inflammation represented by elevated C-reactive protein (CRP) and IFN signatures (David and Frémond, 2022; Frémond et al., 2021; Landman et al., 2020; Wan et al., 2022). To date, more than one dozen MITA GOFs have been identified in the hinge region [V147L/M (Abid et al., 2020; Munoz et al., 2015), N154S and V155M (Jeremiah et al., 2014; Liu et al., 2014), F153V and G158A (Lin et al., 2021)] and the LBD [C206Y/G (Melki et al., 2017), G207E (Keskitalo et al., 2019), F269S (Valeri et al., 2024), F279L (Seo et al., 2017), R281Q/W (Melki et al., 2017; Volpi et al., 2019), and R284S (Konno et al., 2018; Melki et al., 2017; Saldanha et al., 2018)], all of which are localized at the Golgi apparatus and exhibit constitutive polymerization and activation. For example, mutations at F269 or R281 destabilize the side-by-side and head-to-head autoinhibitory conformation of MITA (Liu et al., 2023), whereas mutations at R284 result in the release of the autoinhibitory CTT domain and make the polymerization interface of MITA constitutively available (Ergun et al., 2019). MITA GOFs at the hinge region, such as N154S and V155M, are the most common mutations for SAVI. It is proposed that mutations at the hinge region of MITA disrupt the crossover structure and thereby mimic the binding of cGAMP to the MITA dimer for constitutive ER-to-Golgi transport (Valeri et al., 2024). However, how the spontaneous activation of MITA hinge-region GOFs is regulated remains incompletely understood and lacks direct structural evidence due to the difficulty in sample preparation of these GOFs.

Given the elevated type I IFN signatures in SAVI patients, Janus kinase (JAK) inhibitors such as ruxolitinib, baricitinib (JAK1/2 inhibitors), and tofacitinib (JAK1/3 inhibitor) that downregulate type I IFN signaling have been applied to SAVI patients (Dai et al., 2020; David and Frémond, 2022). However, the results from the available clinical trials have shown large variations in efficacy, with some cases of remission in skin and respiratory symptoms and systemic inflammation (Cetin Gedik et al., 2022; Kanazawa et al., 2023; Li et al., 2022; Sanchez et al., 2018; Seo et al., 2017; Tokgun et al., 2023), and others of limited or no benefit from such treatments (Frémond et al., 2021; Tang et al., 2020; Wan et al., 2022; Wang et al., 2021). The reason behind this phenomenon is unclear but might involve the different MITA GOFs in the reported cases and the ages at the start of treatment. In this context, the clinical benefit of ruxolitinib in SAVI patients is associated with treatment early in the disease course (Frémond et al., 2021), while it is noted that treatment with JAK inhibitors at an early age increases the risk of severe infections, such as shingles, viral respiratory infections, rotavirus enteritidis, and aspergilloma in the lung cavity (Balci et al., 2020; Sanchez et al., 2018; Volpi et al., 2019; Wang et al., 2021). Therefore, elucidating the activation mechanisms of MITA GOFs would not only advance our understanding of the progression of SAVI but also provide precise therapeutic medication for the treatment of SAVI.

iRhom2 (also known as RHBDF2) is an inactive member of the multi-span transmembrane Rhomboid family proteins and exhibits various functions in lupus nephritis (Qing et al., 2018), innate antiviral immunity (Luo et al., 2017, 2016), and cancer (Al-Salihi and Lang, 2020; Xu et al., 2020). Here, we found that the MITA GOFs constitutively interacted with iRhom2 for ER-to-Golgi translocation to promote the progression of SAVI and that knockout of iRhom2 substantially abrogated the SAVI phenotypes caused by MITA^N153S^. In addition, an additional mutation at Lys150 on human MITA^N154S^ (mMITA^N153S^) significantly impaired its association with iRhom2 and the subsequent ER-to-Golgi translocation and failed to induce the expression of type I IFNs and proinflammatory cytokines. Consistently, the SAVI phenotypes, including autoimmune lethality, systemic inflammation, T cell cytopenia and myeloid cell expansion in MITA^N153S/WT^ mice, were completely abolished in MITA^K150N/N153S^ (MITA^NS/NS^) mice and *iRhom2*^-/-^ MITA^N153S/WT^→WT chimeric mice. Interestingly, however, cGAMP could bind to MITA^K150N^ and MITA^NS^ and induce the MITA^NS^-iRhom2 interaction and the ER-to-Golgi translocation of MITA^NS^, indicating that Lys150 is dispensable for cGAMP-induced activation of MITA but required for the activation of MITA hinge-region GOFs. Accordingly, we designed a SAVI-inhibitory peptide (SIP) that selectively impaired the association between MITA hinge-region GOFs and iRhom2 and the expression of type I IFNs and proinflammatory cytokines downstream of MITA hinge-region GOFs but had a minimal effect on cGAMP-induced MITA-iRhom2 interaction and expression of downstream genes. Moreover, treatment with SIP almost completely abrogated the SAVI phenotypes in the MITA^N153S/WT^→WT chimeric mice. These findings have uncovered a previously uncharacterized mechanism for the activation of MITA GOFs and highlighted a potential therapeutic strategy for SAVI.

## 2. Results

### 2.1. Lys150 of MITA hinge-region GOFs is required for their spontaneous activation

It has been recognized that SAVI is caused by gain-of-function mutations (GOFs) of MITA (Jeremiah et al., 2014; Liu et al., 2014). To date, more than one dozen MITA GOFs have been identified, and approximately 80% have been found in the hinge region (V147L, N154S, and V155M) (Frémond et al., 2021; Wang et al., 2021) that contains one lysine residue (Lys150) (Figure S1A). We have previously demonstrated that mutation of Lys150 into Arg (MITA^K150R^) does not affect its ability to activate the IFN-stimulated responsive element (ISRE) in reporter assays (Zhong et al., 2009). Interestingly, introducing a mutation at Lys150 in a MITA hinge-region GOF (MITA^N154S^), such as polar uncharged N, S or Y, non-polar V or F, polar positively charged R, or negatively charged D, substantially impaired MITA^N154S^-mediated activation of the IFN-β promoter in reporter assays (Figure S1B). In addition, mutation of Lys150 into Asn in MITA^V147L^ or MITA^V155M^ significantly inhibited their ability to activate the IFN-β promoter (Figure S1C). In contrast, however, MITA^C206Y/K150N^, MITA^R281Q/K150N^ or MITA^R284S/K150N^ activated the IFN-β promoter to a similar extent as MITA^C206Y^, MITA^R281Q^ or MITA^R284S^, respectively (Figure S1C).

Analyses of mouse MITA suggested that two lysine residues (Lys150 and Lys151) exist in the hinge region of mMITA (Figure S1A). Interestingly, mutation of Lys150 but not Lys151 into Asn resulted in the functional loss of mMITA^N153S^ regarding the capability of activating the IFN-β promoter (Figure S1D-E). In addition, mMITA^V146L/K150N^ and mMITA^V154M/K150N^ failed to activate the IFN-β promoter as did mMITA^V146L^ and mMITA^V154M^, whereas mMITA^C205Y/K150N^, mMITA^R280Q/K150N^, and mMITA^R283S/K150N^ activated the IFN-β promoter to a similar extent as mMITA^C205Y^, mMITA^R280Q^, and mMITA^R283S^, respectively (Figure S1E). These data suggest that Lys150 is conserved and essential for the activity of MITA hinge-region GOFs.

Consistent with the results obtained in the reporter assays, reconstitution of mMITA^N153S^ but not mMITA^K150N/N153S^ (mMITA^NS^) into *Mita*^-/-^ mouse lung fibroblasts (MLFs) induced the expression of *Ifnb*, *Cxcl10*, *Isg56* or *Isg15* or triggered the production of CXCL1, TNFα or IL-6 (Figure 1A-B), suggesting that the mutation of Lys150 leads to the inactivation of MITA hinge-region GOFs. In contrast, reconstitution of mMITA^R283S^ or mMITA^R283S/K150N^ into *Mita*^-/-^MLFs similarly upregulated the mRNA levels of *Ifnb*, *Isg15*, *Isg56* and *Cxcl10* (Figure S1F). These data suggest that mutation of Lys150 leads to the inactivation of MITA hinge-region GOFs.

**Figure 1.**
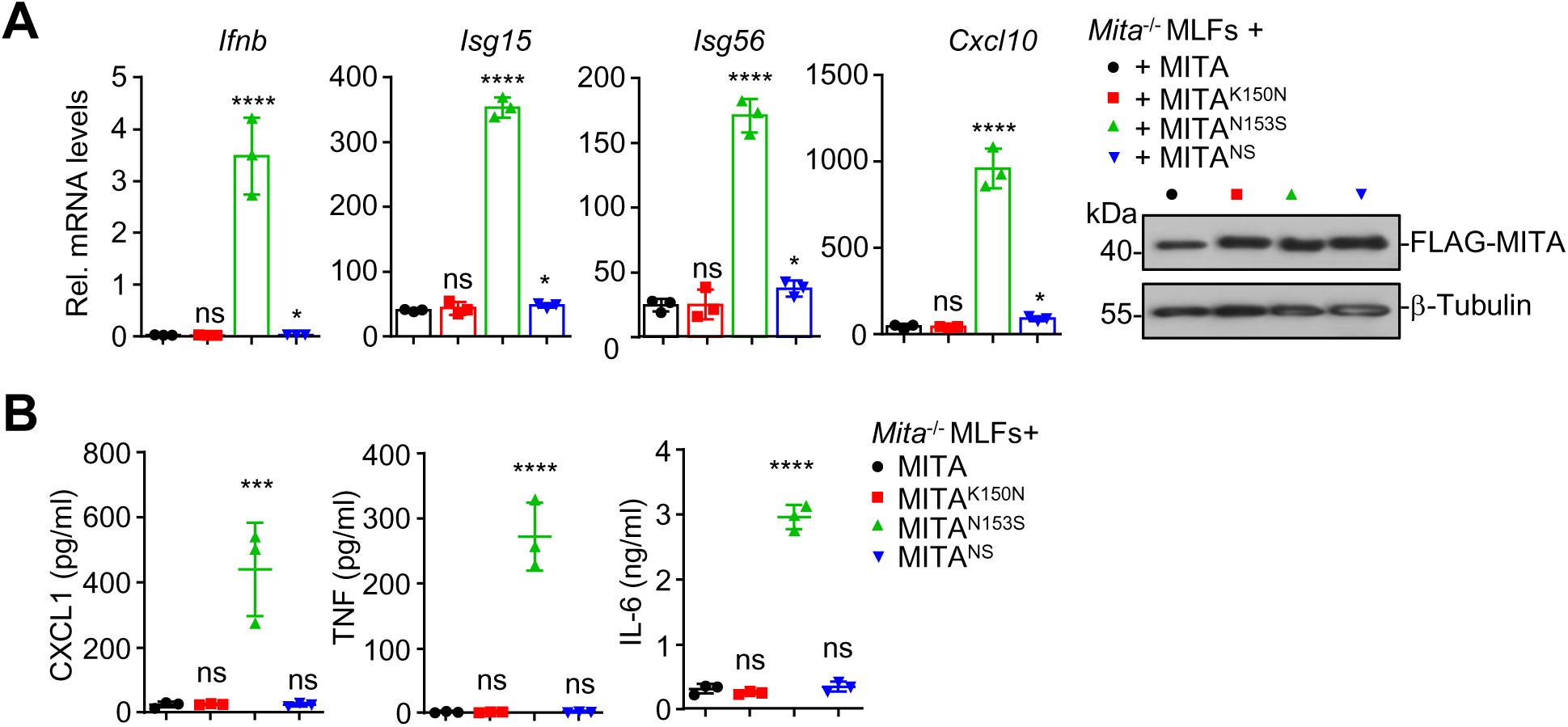
Mutation of K150 impairs the activity of MITA^N153S^. (A) RT-qPCR analysis of *Ifnb*, *Isg15*, *Isg56*, and *Cxcl10* mRNA (left graphs) and immunoblot analysis of MITA, MITA^K150N^, MITA^N153S^, or MITA^K150N/N153S^ of *Mita*^-/-^ MLFs reconstituted with wild-type (WT) or different MITA mutants. (B) ELISA analysis of CXCL1, TNF, and IL-6 in the supernatants of *Mita*^-/-^ MLFs reconstituted with wild-type or different MITA mutants. \**P* <0.05, \*\*\**P*<0.001, \*\*\*\**P*<0.0001; ns, not significant (one-way ANOVA). The graphs show mean ± SD. The data are representative of three independent experiments.

### 2.2. The SAVI autoimmune phenotypes are completely abolished in MITA^NS/NS^ mice

To further substantiate the role of Lys150 in the activation of MITA hinge-region GOFs, we generated MITA^N153S/WT^ (Zhang et al., 2024) and MITA^K150N/N153S^ (designated MITA^NS/NS^) knock-in mice via CRISPR/Cas9-mediated genome editing (Figure S2A). Unlike the homozygous MITA^N153S/N153S^ mice that are embryonic lethal (Luksch et al., 2019; Motwani et al., 2019; Warner et al., 2017; Zhang et al., 2024), the homozygous MITA^NS/NS^ mice were viable and exhibited normal growth and reproduction compared to the wild-type controls, and the sex ratio of the offspring of the MITA^NS/NS^ breeders conformed to the Mendelian inheritance law (Figure 2A and Figure S2B-C). The protein levels of MITA and MITA mutants were comparable in the kidneys and the hearts of wild-type, MITA^N153S/WT^, and MITA^NS/NS^ mice (Figure S2D). These results suggest that simultaneous mutation of Lys150 into Asn and Asn153 into Ser in MITA does not affect the growth or the development of mice.

**Figure 2.**
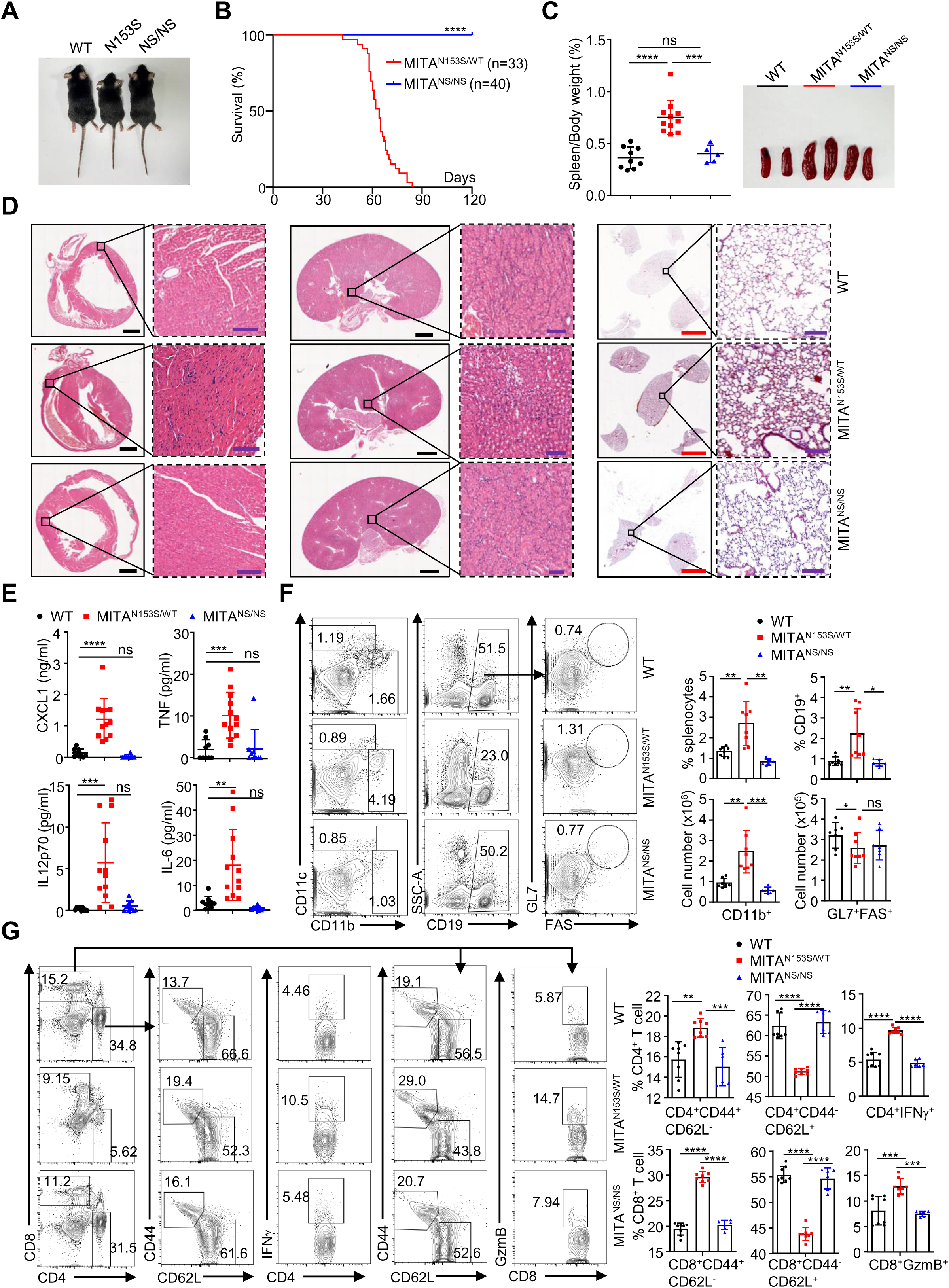
The SAVI phenotypes are abolished in MITA^K150N/N153S^ mice. (A) Representative images of 5-week-old WT, MITA^N153S/WT^ and MITA^NS/NS^ mice. (B) Survival analysis (Kaplan–Meier curves) of MITA^N153S/WT^ (n=33) and MITA^NS/NS^ (n=40) mice. (C) Spleen weight/body weight ratios (left) and representative images (right) of the spleens of 5-week-old WT (n=9), MITA^N153S/WT^ (n=11), and MITA^NS/NS^ (n=5) mice. (D) Representative images of HE-stained heart, kidney, and lung sections from 5-week-old WT (n=3), MITA^N153S/WT^ (n=3), and MITA^NS/NS^ (n=3) mice. (E) ELISA analysis of CXCL1, TNF, IL12p70, and IL-6 in the sera of 5-week-old WT (n=9), MITA^N153S/WT^ (n=12) and MITA^NS/NS^ (n=9) mice. (F-G) Flow cytometric analysis of splenocytes from 5-week-old WT (n=8), MITA^N153S/WT^ (n=8), and MITA^NS/NS^ (n=6) mice stained with fluorophore-conjugated antibodies against the indicated surface or intracellular molecules. \**P* <0.05, \*\**P*<0.01, \*\*\**P*<0.001, \*\*\*\**P*<0.0001; ns, not significant (one-way ANOVA or the log-rank test). Scale bars represent 1 mm (black), 100 μm (purple), or 3 mm (red) in (D). The graphs show the means ± SD. The data are combined results from two independent experiments (B) or representative of three independent experiments (A, C-G)

Importantly, compared to the MITA^N153S/WT^ mice, the MITA^NS/NS^ mice of the same age and sex had a larger body size, heavier body weight, and smoother fur, which were comparable to those of the wild-type mice (Figure 2A and Figure S2C). In addition, the autoimmune lethality was completely rescued in the MITA^NS/NS^ mice, whereas the MITA^N153S/WT^ mice began to die 35 days after birth and had a median survival of 64 days (Zhang et al., 2024) (Figure 2B). We next analyzed the autoimmune SAVI phenotypes in MITA^NS/NS^ mice. First, the spleen sizes were substantially smaller and the spleen weight/body weight ratios were lower in MITA^NS/NS^ mice than in MITA^N153S/WT^ mice (Figure 2C). Second, the results from HE staining revealed substantially fewer infiltrated immune cells in the heart, lung, or kidney of MITA^NS/NS^ mice than in the counterparts of MITA^N153S/WT^ mice (Figure 2D). Third, the results from RT-qPCR analyses suggested that the mRNA levels of *Ccl2*, *Ccl3*,

*Ccl4*, and *Tnf* in the liver, brain, and MLFs of MITA^NS/NS^ mice were significantly lower than those in the liver, brain, and MLFs of MITA^N153S/WT^ mice (Figure S2E). In addition, the protein levels of CXCL1, TNFα, IL-12p70, and IL-6 in the serum of MITA^NS/NS^ mice were significantly lower than those in the serum of MITA^N153S/WT^ mice (Figure 2E). Finally, compared to those of MITA^N153S/WT^ mice, the proportions and numbers of Lin^-^Sca1^-^c-kit^+^ myeloid progenitors in the bone marrow, CD4^-^ CD8^-^ thymocytes in the thymus, CD3^+^ T cells in the blood, and CD4^+^ and CD8^+^ T cells and CD19^+^ B cells in the spleen of the MITA^NS/NS^ mice were significantly increased, whereas the proportions and numbers of CD11b^+^ myeloid cells in blood and spleen and FAS^+^GL7^+^ germinal center B cells (GCBs) in spleen and the proportions of CD4^+^IFNγ^+^ T cells and CD8^+^GzmB^+^ T cells in spleen of the MITA^NS/NS^ mice were significantly decreased (Figure 2F-G and Figure S2F-I). Notably, all of the examined autoimmune SAVI phenotypes, including developmental retardation, systemic inflammation, and dysregulated development of immune cells, were abrogated in the MITA^NS/NS^ mice and were similar to those of the wild-type controls. These data demonstrate that Lys150 of MITA critically coordinates the SAVI autoimmunity caused by MITA^N153S^.

### 2.3. MITA^NS^ is localized on the ER and transported to the Golgi apparatus by cGAMP

MITA GOFs such as MITA^N153S^ are constitutively and spontaneously transported from the endoplasmic reticulum (ER) to the Golgi apparatus where they activate downstream signaling pathways for SAVI progression (Dobbs et al., 2015; Jeremiah et al., 2014; Zhang et al., 2020) (Figure 3A). In contrast, MITA^NS^ was located at the ER as wild-type MITA or MITA^K150N^ (Figure 3A), and cGAMP treatment resulted in the translocation of MITA^NS^ and MITA^K150N^ from the ER to the Golgi apparatus similarly as wild-type MITA (Figure 3A).

**Figure 3.**
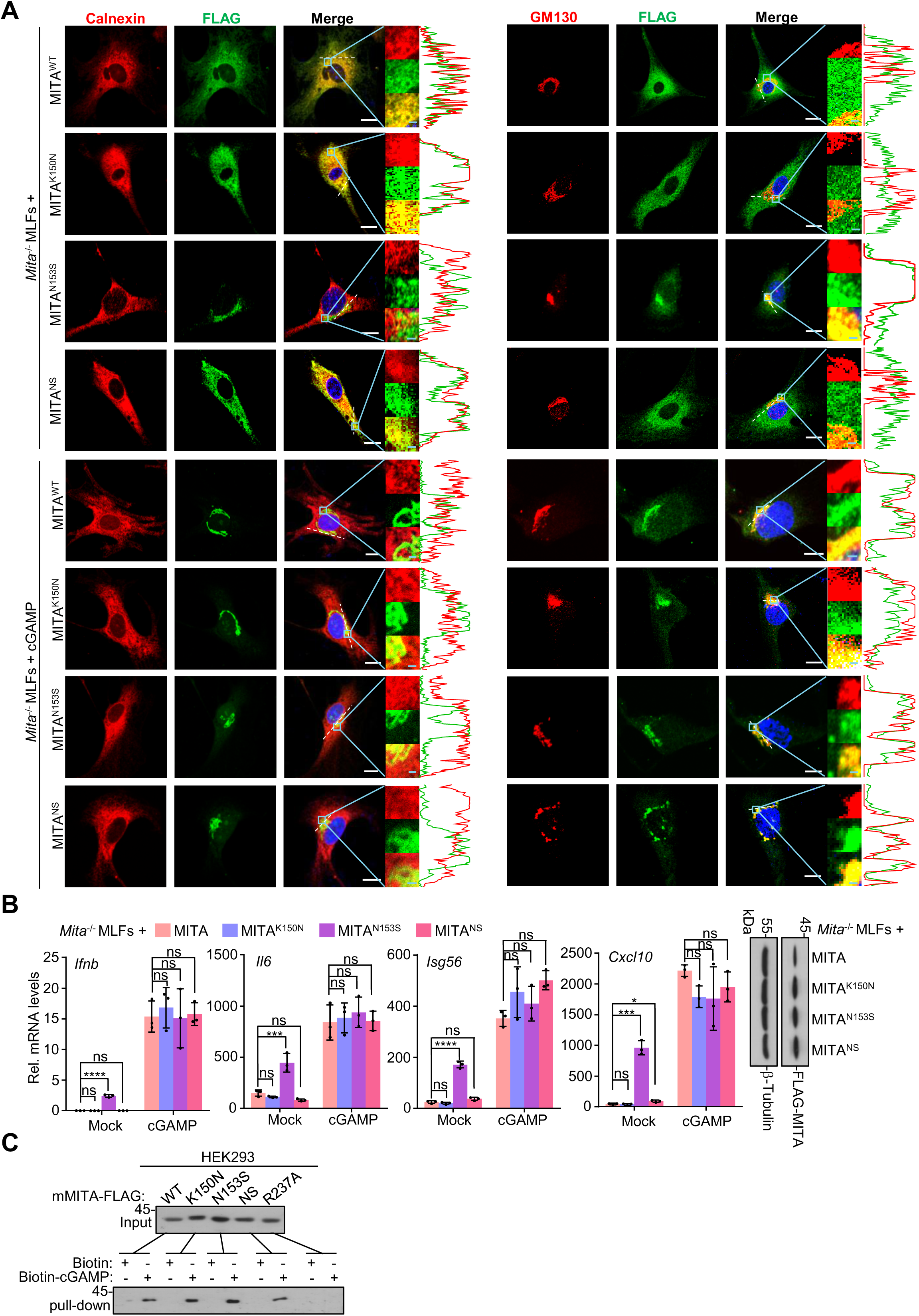
MITA^NS^ binds and responds to cGAMP. (A) Immunofluorescence staining of Calnexin (an ER marker) (red), GM130 (a Golgi apparatus marker) (red), and FLAG (green) and confocal microscopy analysis of *Mita*^-/-^ MLFs reconstituted with FLAG-tagged MITA, MITA^K150N^, MITA^N153S^ or MITA^NS^ and treated with or without cGAMP (1 μg/mL) for 4 h, and co-localization analysis was conducted. (B) RT-qPCR analysis of *Ifnb*, *Il6*, *Isg56*, and *Cxcl10* mRNA (left graphs) and immunoblot analysis of MITA, MITA^K150N^, MITA^N153S^, or MITA^NS^ of *Mita ^-/-^* MLFs reconstituted with FLAG-tagged MITA^WT^, MITA^K150N^, MITA^N153S^, or MITA^NS^ followed by treatment with cGAMP (1 μg/mL) for 0-4 h. (C) cGAMP binding assays of MITA^WT^, MITA^K150N^, MITA^N153S^, or MITA^NS^ that were transfected into HEK293 cells followed by in vitro pulldown by Biotin-cGAMP. \**P* <0.05, \*\*\**P*<0.001, \*\*\*\**P*<0.0001; ns, not significant (one-way ANOVA). Scale bars represent 10 μm (white) or 1 μm (cyan). The graphs show mean ± SD. The data are representative of two independent experiments.

Consistent with the results of the immunofluorescent staining analysis, cGAMP treatment significantly increased the mRNA levels of *Ifnb*, *Il6*, *Isg56*, and *Cxcl10* in *Mita*^-/-^ MLFs reconstituted with MITA^NS^ to a similar extent as those in *Mita*^-/-^ MLFs reconstituted with wild-type MITA or MITA^K150N^ (Figure 3B). In addition, similar to wild-type MITA or MITA^K150N^, MITA^NS^ bound to cGAMP in *in vitro* cGAMP pulldown assays (Figure 3C). Taken together, these data suggest that Lys150 selectively contributes to the activity of MITA hinge-region GOFs but not the cGAMP-induced activation of MITA.

### 2.4. MITA^NS^ adopts conformations similar to wild-type MITA in both apo and cGAMP-bound states

We next used cryoelectron microscopy (cryo-EM) to determine the structure of apo-hMITA^NS^ expressed in HEK293F cells and obtained the 3D structure at a resolution of 3.9 Å (Figure S3 and Table S1). Similar to wild-type MITA (Shang et al., 2019), the apo-MITA^NS^ structure adopted a domain-swapped dimer conformation with a crossed hinge region (Figure 4A), confirming that apo-MITA^NS^ is inactive. In wild-type MITA, the residue N154 formed a hydrogen bond with the residue N154′ on the opposing protomer (Figure 4B), which is responsible for the stabilization of the crossover structure and the inactive state of MITA dimer. However, MITA^NS^ dimer that lost the hydrogen bond between N154 and N154′ still maintained the crossover structure and remained in an inactive state (Figure 4B), implicating that K150 facilitates N154S mutation-induced destabilization of the MITA dimer.

**Figure 4.**
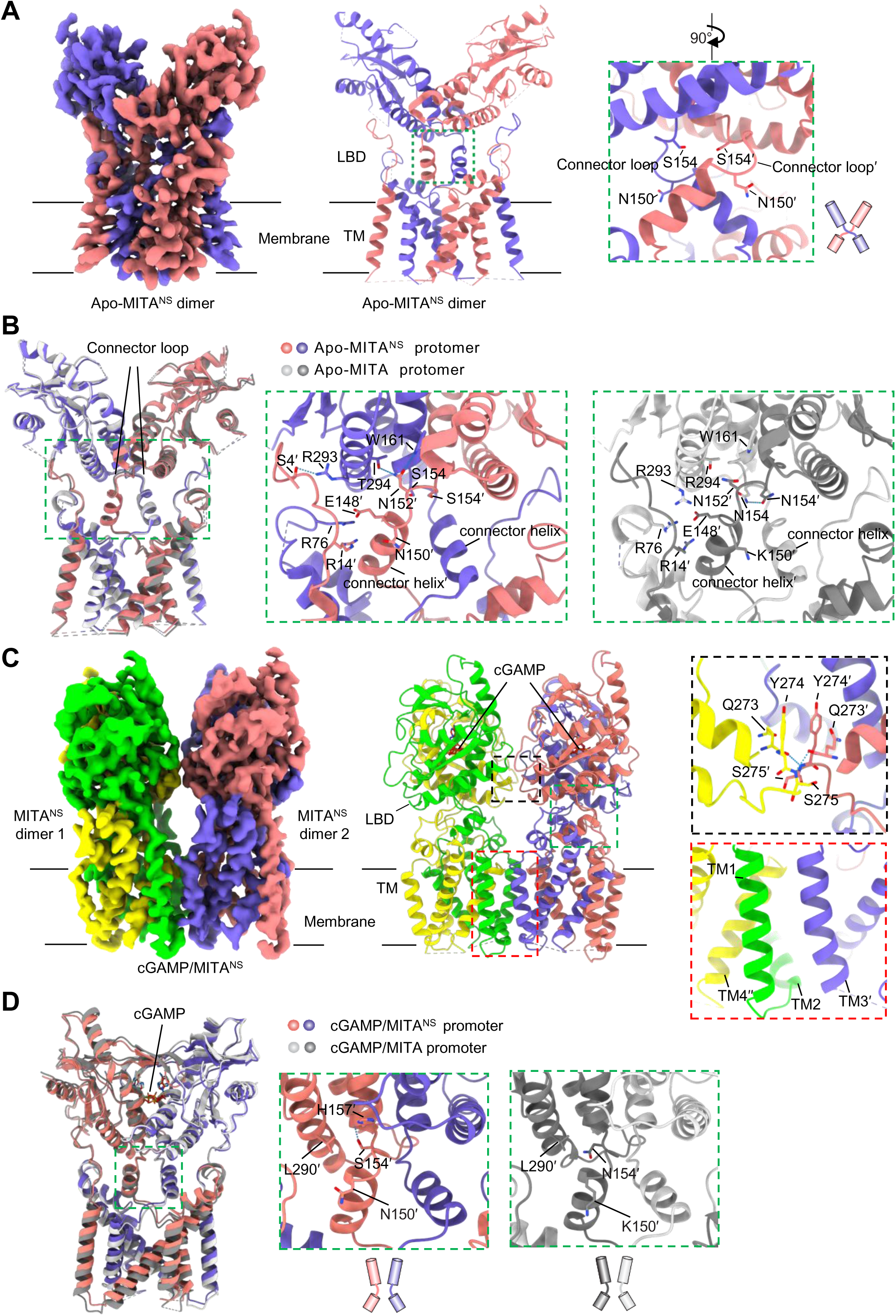
Structural analyses of apo-MITA^NS^ and cGAMP/MITA^NS^ complex. (A) Cryo-EM map of apo-MITA^NS^ (left), and cartoon representation showing domain-swapped dimer conformation with crossed hinge region (middle), and the hinge region (right) of the MITA^NS^ dimer. (B) Structural comparison between wild-type apo-MITA (PDB: 6NT5) and apo-MITA^NS^ (PDB: 9VXT). The shift of connector loop in apo-MITA^NS^ relative to it in wild-type apo-MITA is indicated by red arrow. Inset: the residues located around hinge region were shown (middle panel: apo-MITA^NS^; right panel: wild-type apo-MITA). (C) Cryo-EM map of cGAMP/MITA^NS^ filament (tetramer shown) (upper left) (PDB ID: 9VXU), cartoon presentation of cGAMP/MITA^NS^ filament (tetramer shown) (upper right), and interactions between LBDs (lower left) and TM (lower middle) from adjacent cGAMP/MITA^NS^ dimer and interaction network in the hinge region of cGAMP/MITA^NS^ dimer (lower right). The scheme on the right shows the connector region in a parallel conformation. (D) Structural comparison between cGAMP/MITA dimer (PDB: 8IK3) and cGAMP/MITA^NS^ (PDB: 9VXU). Inset: the residues located around the hinge region were shown (middle panel: cGAMP/MITA^NS^; right panel: cGAMP/MITA). TM: transmembrane domain; LBD: ligand-binding domain.

Interestingly, we observed that the side chain of N152 formed two new hydrogen bonds with W161 and T294 of the opposing ligand-binding domain (LBD) (Figure 4B). Furthermore, while the conserved residue E148 engaged in charge-charge interactions with R14′, R76, and R293 in the wild-type apo-MITA structure, it interacted only with R76 in apo-MITA^NS^ and R293 reoriented to form an interaction with residue S4′ on the opposite protomer.

Concomitantly, the connector loop of the hinge region shifted outward relative to its position in wild-type apo-MITA (Figure 4B). Collectively, these newly established features might contribute to the stabilization of the inactive conformation of apo-MITA^NS^.

To elucidate how MITA^NS^ responded to cGAMP as did wild-type MITA, we determined the structure of cGAMP-bound MITA^NS^ by Cryo-EM and obtained a polymerized cGAMP/MITA complex at a resolution of 3.2 Å (Figure S4 and Table S1). The cGAMP-bound MITA^NS^ polymer formed a curved filament stabilized by side packing of MITA^NS^ dimers (Figure 4C). Each dimer within the polymer assumed a closed conformation, with cGAMP buried within the ligand-binding pocket beneath the four-stranded lid (Figure 4C). The connector helix adopted a parallel orientation, and inter-dimer interactions were mediated by both the LBD and the TM (Figure 4C). Specifically, LBD interactions involved hydrogen bonding between the main chains of adjacent LBDα2-α3 loops (residues Q273, Y274, and S275) and TM interactions involved TM1 from one dimer packing against TM2-TM3′ and TM4′′ from an adjacent dimer (Figure 4C), mirroring the key features of the cGAMP/MITA complex (Liu et al., 2023). We also noted a difference at residue 154 between cGAMP-bound MITA^NS^ and cGAMP-bound MITA. The S154 residue in cGAMP-bound MITA^NS^ formed an intramolecular hydrogen bond with H157, whereas the side chain of N154 in cGAMP-bound MITA was engaged in interaction with L290 (Figure 4D). Nonetheless, as evidenced by the low root mean square deviation (RMSD) of 1.3 Å for the aligned Cα atoms (Figure 4D), the overall structure of cGAMP/MITA^NS^ complex is similar to that of the cGAMP/MITA complex, implying similar activation mechanisms of MITA^NS^ and wild-type MITA by cGAMP.

### 2.5. Mutation at Lys150 in MITA GOFs impairs their interaction with iRhom2

The above apo-MITA^NS^ structure implicated a critical role of K150 in the destabilization of the inactive state of MITA^N154S^ (Figure 4A-B). Together with the observations that mMITA^NS^ was localized on the ER and mMITA^N153S^ was localized on the Golgi apparatus (Figure 3A), we reasoned that the mutation at Lys150 might affect the interaction between MITA^N154S^ and ER retention factors or ER-to-Golgi trafficking factors for destabilizing the inactive state and trafficking from the ER to the Golgi apparatus. To test this hypothesis, we performed screening assays by transfecting FLAG-MITA^N154S^ or MITA^NS^ together with HA-tagged retention factor (STIM1) (Srikanth et al., 2019) or trafficking factors [EXOC2 (Ishikawa et al., 2009), SAR1a (Zhang et al., 2020), iRhom2 (Luo et al., 2016), SURF4 (Mukai et al., 2021), STEEP (Zhang et al., 2020), Sec24c (Ishikawa and Barber, 2008), TRAPβ (Ishikawa and Barber, 2008) and TMED2(Sun et al., 2018)] into HEK293 cells followed by co-immunoprecipitation analysis (Figure S5A-B). The results suggested that the MITA^NS^-iRhom2 and the MITA^NS^-SURF4 associations were substantially weaker than the MITA^N154S^-iRhom2 and the MITA^N154S^-SURF4 associations, respectively (Figure S5A).

Although SURF4 serves as a cargo receptor that facilitates both ER-to-Golgi and retrograde Golgi-to-ER transport of cargos (Shen et al., 2023), a recent study has shown that SURF4 interacts with the coat protein complex I (COPI) α subunit (α-COP) to facilitate Golgi-to-ER transport of MITA and negatively regulate the activation of MITA (Mukai et al., 2021).

Considering that SURF4 is localized mainly within the ER-Golgi intermediate compartment (ERGIC) and early Golgi apparatus (Mitrovic et al., 2008), the increased MITA^N154S^-SURF4 associations versus MITA^NS^-SURF4 associations were likely due to the Golgi localization of MITA^N154S^ versus the ER localization of MITA^NS^ (Figure 3A). We have previously demonstrated that iRhom2 is localized on the ER and translocated to the ERGIC after HSV-1 infection and promotes the ER-to-Golgi trafficking and activation of MITA (Luo et al., 2016). Interestingly, we further found that the associations between MITA^V147L/K150N^ or MITA^V155M/K150N^ and iRhom2 were substantially weaker than those between MITA^V147L^ or MITA^V155M^ and iRhom2, respectively (Figure S5C), and mMITA^NS^ exhibited decreased interaction with iRhom2 compared to mMITA^N153S^ (Figure S5D-E). In contrast, however, the associations between MITA^C206Y/K150N^, MITA^R281Q/K150N^, or MITA^R284S/K150N^ and iRhom2 were comparable to those between MITA^C206Y^, MITA^R281Q^, or MITA^R284S^ and iRhom2 (Figure S5C), indicating that Lys150 specifically regulates the associations between iRhom2 and MITA hinge-region GOFs. In this context, our earlier findings revealed that a mutation at Lys150 selectively impaired the activity of MITA hinge-region GOFs (Figure S1C and E).

Mouse MITA^NS^ is translocated from the ER to the Golgi apparatus upon cGAMP treatment to induce the expression of downstream genes (Figure 3A-B). Interestingly, we found that cGAMP treatment induced the association between MITA^NS^ and iRhom2 (Figure S5F), and that knockdown of iRhom2 significantly inhibited cGAMP-induced upregulation of downstream genes such as *Ifnb*, *Il6*, *Cxcl10*, *Isg56* and *Isg15* in *Mita*^NS/NS^ MLFs (Figure S5G), indicating that MITA^NS^ responds to cGAMP for downstream signaling in a manner dependent on iRhom2. These data align with our previous observations that the structure of cGAMP/MITA^NS^ complex resembles that of cGAMP/MITA and that cGAMP treatment induces the interaction between MITA and iRhom2 (Liu et al., 2023; Luo et al., 2016) (Figure 4C and Figure S4). Collectively, these data suggest that Lys150 is selectively required for the optimal association between MITA hinge-region GOFs and iRhom2 but not for the association between cGAMP-bound MITA and iRhom2.

### 2.6. iRhom2 is required for the activation of MITA^N153S^ and the progression of SAVI

We next examined whether iRhom2 was required for the activation of hMITA^N154S^ and mMITA^N153S^. As shown in Figure S6A, knockdown of iRhom2 inhibited hMITA^N154S^-mediated activation of the IFN-β promoter in HEK293 cells in reporter assays. The expression of *Ifnb*, *Il6*, *Cxcl10* and *Isg56* was significantly downregulated in *iRhom2*^−/−^MITA^N153S/WT^ MLFs compared to MITA^N153S/WT^ MLFs (Figure S6B), suggesting an indispensable role of iRhom2 for the activation of MITA^N153S^. We next generated bone marrow chimeric mice by adoptively transferring bone marrow cells from 5-week-old MITA^N153S/WT^ or *iRhom2*^−/−^MITA^N153S/WT^ mice into irradiated wild-type C57BL/6 mice, and observed that the *iRhom2^−/−^*MITA^N153S/WT^→WT chimeric mice exhibited significantly increased survival time and weight gain and reduced spleen sizes compared to the MITA^N153S/WT^→WT chimeric mice (Figure 5A-C). The systemic inflammation, including immune cell infiltration, the mRNA levels of proinflammatory cytokines in different organs and the protein levels of proinflammatory cytokines in the blood, was significantly compromised in *iRhom2^−/−^*MITA^N153S/WT^→WT chimeric mice compared to MITA^N153S/WT^→ WT chimeric mice (Figure 5D-E and Figure S6C). In addition, the dysregulated development of immune cells in the bone marrow (Lin^-^c-Kit^+^ myeloid progenitors), thymus (CD4^-^CD8^-^thymocytes), blood (CD11b^+^F4/80^+^ macrophages and CD11b^+^Ly6G^+^ neutrophils) and spleen (CD3^+^, CD4^+^ and CD8^+^ T cells, FAS^+^GL7^+^ GCBs, CD4^+^IFNγ^+^ T cells, CD4^+^CD44^+^CD62L^-^ effector T cells, and CD8^+^GzmB^+^ T cells) was also restored in the *iRhom2^−/−^*MITA^N153S/WT^→ WT chimeric mice compared to the MITA^N153S/WT^→WT chimeric mice (Figure 5F-G and Figure S6D-F). Taken together, these data demonstrate that iRhom2 is required for MITA^N153S^-induced SAVI autoimmunity in mice.

**Figure 5.**
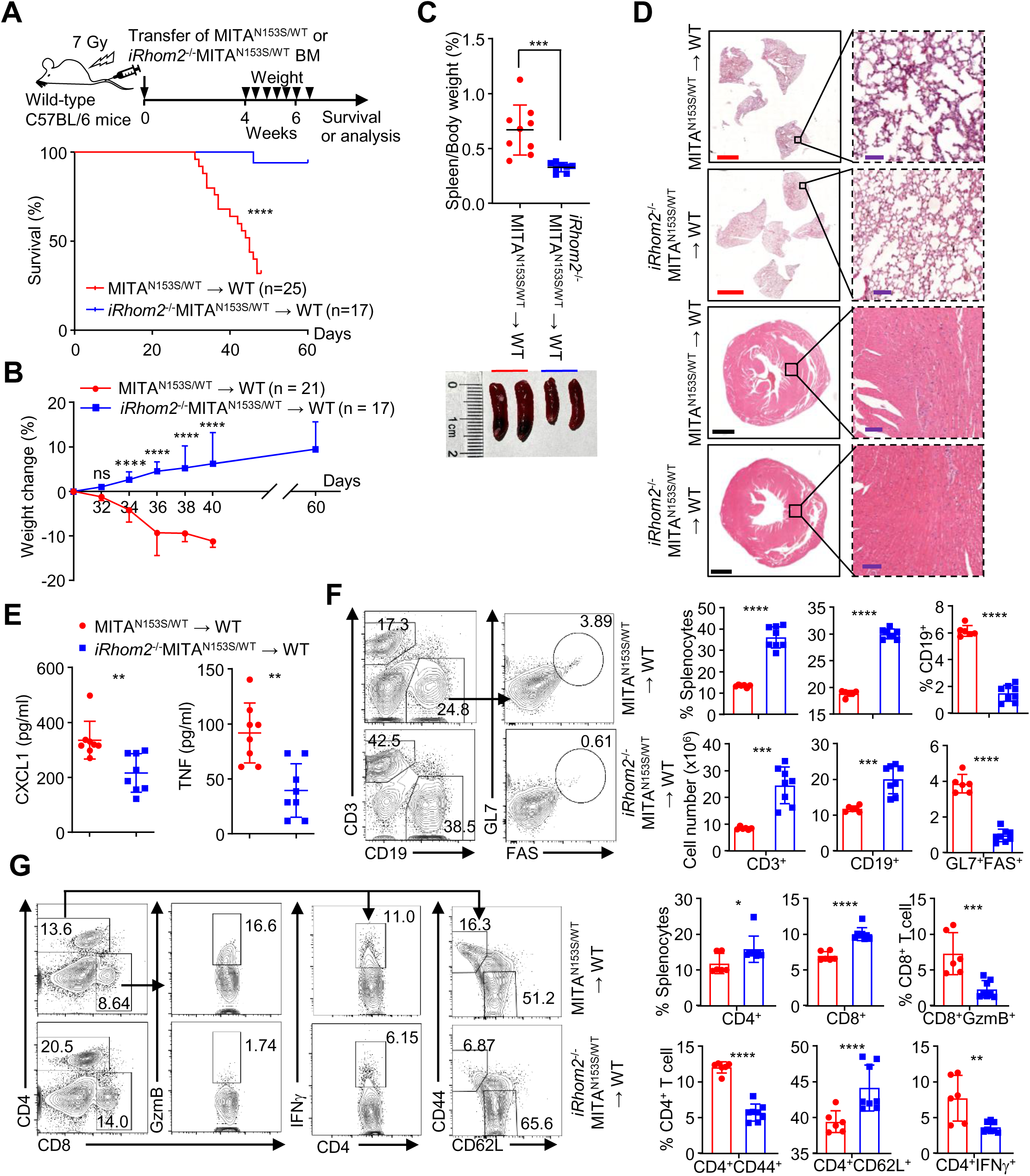
iRhom2 promotes SAVI progression in MITA^N153S/WT^ mice. (A) Survival (Kaplan-Meier curves) of MITA^N153S/WT^→WT (n=25) and *iRhom2^-/-^* MITA^N153S/WT^→WT (n=17) bone marrow chimeric mice. (B) Weight changes of MITA^N153S/WT^→WT (n=21) and *iRhom2^-/-^*MITA^N153S/WT^→WT (n=17) chimeric mice. (C) Spleen weight/body weight ratios (upper) and representative images (lower) of spleens from MITA^N153S/WT^→WT (n=9) and *iRhom2^-/-^*MITA^N153S/WT^→WT (n=8) chimeric mice at the 7^th^ week after bone marrow transfer. (D) Representative images of HE-stained heart and lung sections of MITA^N153S/WT^ →WT (n=3) and *iRhom2^-/-^*MITA^N153S/WT^→WT (n=3) chimeric mice at the 7^th^ week after bone marrow transfer. (E) ELISA of CXCL1 and TNF in the sera of MITA^N153S/WT^→WT (n=8) and *iRhom2^-/-^* MITA^N153S/WT^→WT (n=8) chimeric mice at the 7^th^ week after bone marrow transfer. (F-G) Flow cytometric analysis of splenocytes from MITA^N153S/WT^→WT (n=6) and *iRhom2^-/-^* MITA^N153S/WT^→WT (n=8) chimeric mice at the 7^th^ week after bone marrow transfer. \**P* <0.05, \*\**P*<0.01, \*\*\**P*<0.001, \*\*\*\**P*<0.0001; ns, not significant (one-way ANOVA, the two-tailed Student’s t-test). Scale bars represent 1 mm (black), 100 μm (purple), or 3 mm (red) in (D). The graphs show mean ± SD. The data are combined results from two independent experiments (A-B) or representative of two independent experiments (C-G).

### 2.7. iRhom2 mediates the Golgi localization of MITA GOFs

It has been shown that MITA^N153S^ is constitutively localized on the Golgi apparatus (Dobbs et al., 2015; Zhang et al., 2020) (Figure 3A). Interestingly, we found that knockout of iRhom2 resulted in substantially decreased Golgi localization and increased ER localization of MITA hinge-region GOFs including MITA^V146L^, MITA^N153S^ and MITA^V154M^ (Figure 6A), suggesting that iRhom2 is required for the ER-to-Golgi trafficking of MITA hinge-region GOFs. In addition, similar to our previous observations that iRhom2 promotes HSV-1- or cGAMP-induced ER-to-Golgi translocation of MITA (Luo et al., 2016), cGAMP treatment induced the ER-to-Golgi translocation of MITA^NS^ in *iRhom2*^+/+^ MLFs but not in *iRhom2*^-/-^MLFs (Figure 6B), indicating that iRhom2 is also required for the ER-to-Golgi transportation of cGAMP-bound MITA^NS^. Collectively, these data suggest distinct activation mechanisms for cGAMP-bound MITA and MITA hinge-region GOFs, the former of which is iRhom2-dependent but independent of Lys150 and the latter of which is both iRhom2- and Lys150-dependent.

**Figure 6.**
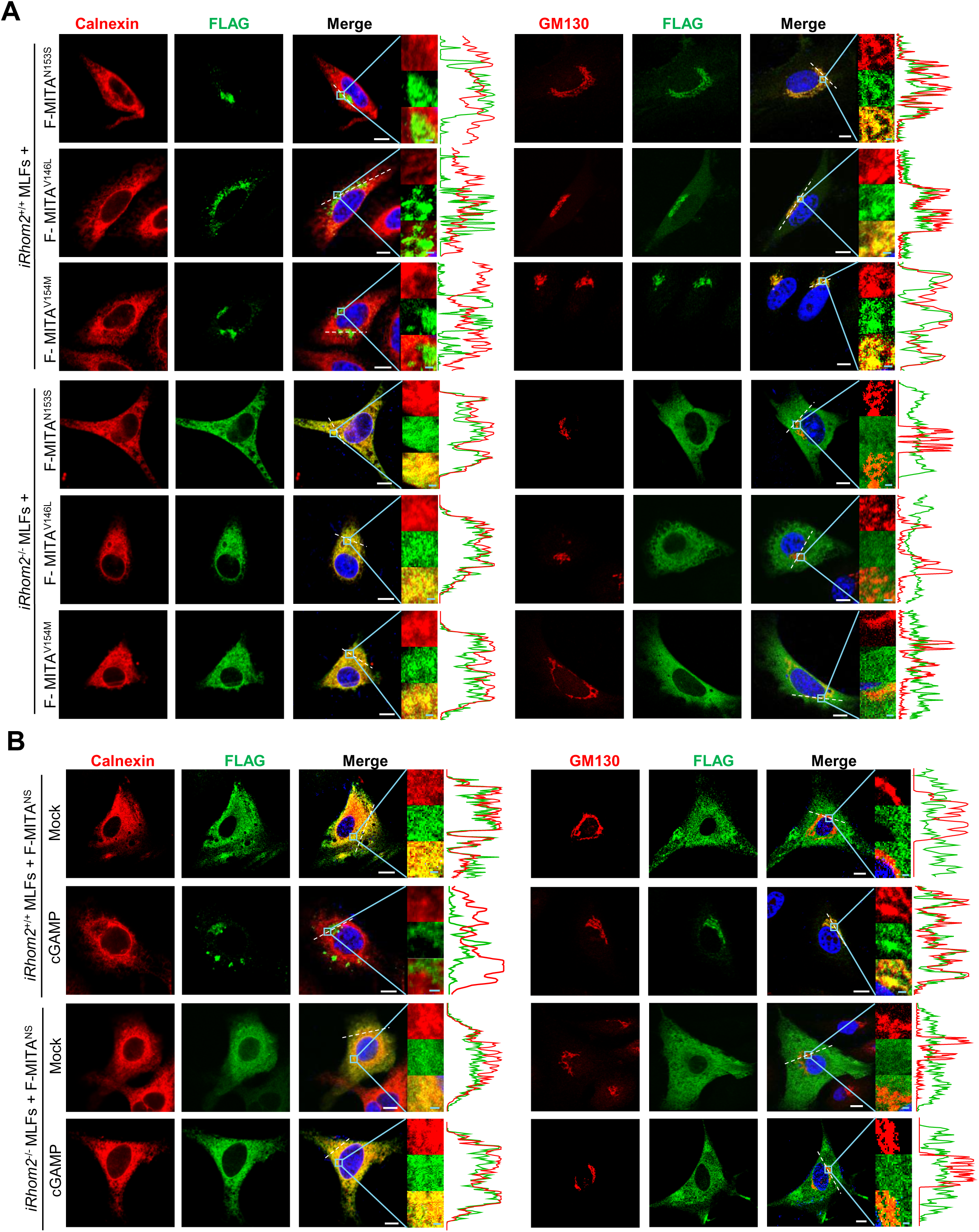
iRhom2 is required for the Golgi localization of MITA GOFs. (A) Immunofluorescence staining of Calnexin (an ER marker) (red), GM130 (a Golgi apparatus marker) (red), and FLAG (green) and confocal microscopy analysis of *iRhom2^+/+^* or *iRhom2^-/-^* MLFs reconstituted with FLAG-tagged MITA^N153S^, MITA^V146L^, or MITA^V154M^, and co-localization analysis was conducted. (B) Immunofluorescence staining of Calnexin (an ER marker) (red), GM130 (a Golgi apparatus marker) (red), and FLAG (green) and confocal microscopy analysis of *iRhom2^+/+^* or *iRhom2^-/-^* MLFs reconstituted with FLAG-MITA^NS^ followed by transfection with cGAMP (1 μg/mL) for 0-4 h, and co-localization analysis was conducted. Scale bars represent 10 μm (white) or 1 μm (cyan). The data are representative of two independent experiments.

### 2.8. The SAVI-inhibitory peptide (SIP) selectively impairs the activation of MITA GOFs

The distinct activation mechanisms of cGAMP-bound MITA and MITA GOFs prompted us to speculate a therapeutic window for SAVI through selectively attenuating the activity of MITA GOFs without affecting cGAMP-induced activation of MITA. Considering that the MITA hinge-region GOFs interacted with iRhom2 dependently on Lys150, we reasoned that peptides spanning the Lys150 and the hinge-region GOFs of MITA might interfere with the iRhom2-MITA hinge-region GOF interactions and thereby impair the activation of MITA GOFs. To test this idea, we synthesized a series of peptides consisting of the transactivator of transcription peptide (TAT), an intermediate “GSG” linker and the wild-type (“CEEKKL<u>N</u>VAH”, TAT-WT) or mutated (“VCEEKKL<u>S</u>VAHGLAWSYYI”, TAT-SIP1; “CEEKKL<u>S</u>VAH”, TAT-SIP2; “CEE<u>N</u>KL<u>S</u>VAH”, TAT-NS) hinge region of MITA (Figure S7A). The results from reporter assays suggested that TAT-SIP1 and TAT-SIP2 but not TAT-WT or TAT-NS significantly inhibited mMITA^N153S^-mediated activation of the IFN-β promoter in HEK293 cells without obvious cytotoxicity (Figure S7A-B). In addition, TAT-SIP2 significantly inhibited mMITA^V146L^ or mMITA^V154M^ - but not mMITA^C205Y^, mMITA^R280Q^, or mMITA^R283S^-mediated activation of IFN-β promoter (Figure S7C). Similarly, the TAT-human SIP2 (hSIP2) peptide spanning Lys150 and the mutated hinge-region GOFs of hMITA (CEKGNF<u>S</u>VAH) significantly inhibited hMITA^V147L^, hMITA^N154S^, or hMITA^V155M^- but not hMITA^C206Y^-, hMITA^R281Q^-, or hMITA^R284S^-mediated activation of the IFN-β promoter (Figure S7D). Consistent with these observations, we found that TAT-SIP1 or TAT-SIP2 inhibited the expression of *Ifnb*, *Ifna4* and *Il12p40* and impaired the associations between iRhom2 and MITA^N153S^ in MITA^N153S/WT^ MLFs in a dose-dependent manner (Figure 7A-B) and that TAT-SIP2 inhibited the upregulation of *Ifnb*, *Isg56* and *Cxcl10* in *Mita*^-/-^ MLFs reconstituted with mMITA^V146L^ or mMITA^V154M^ (Figure S7E). In contrast, TAT-SIP1, TAT-SIP2 or TAT-WT did not inhibit cGAS plus wild-type MITA-induced activation of the IFN-β promoter in HEK293 cells (Figure S7F), impair cGAMP-induced expression of *Ifnb*, *Cxcl10*, *Il6* or *Isg56* (Figure S7G), or impede iRhom2-MITA associations in *Mita*^+/+^ MLFs after HSV-1 infection (Figure S7H). Collectively, these data suggest that the designed SIP peptides selectively inhibit the activation of MITA hinge-region GOFs.

**Figure 7.**
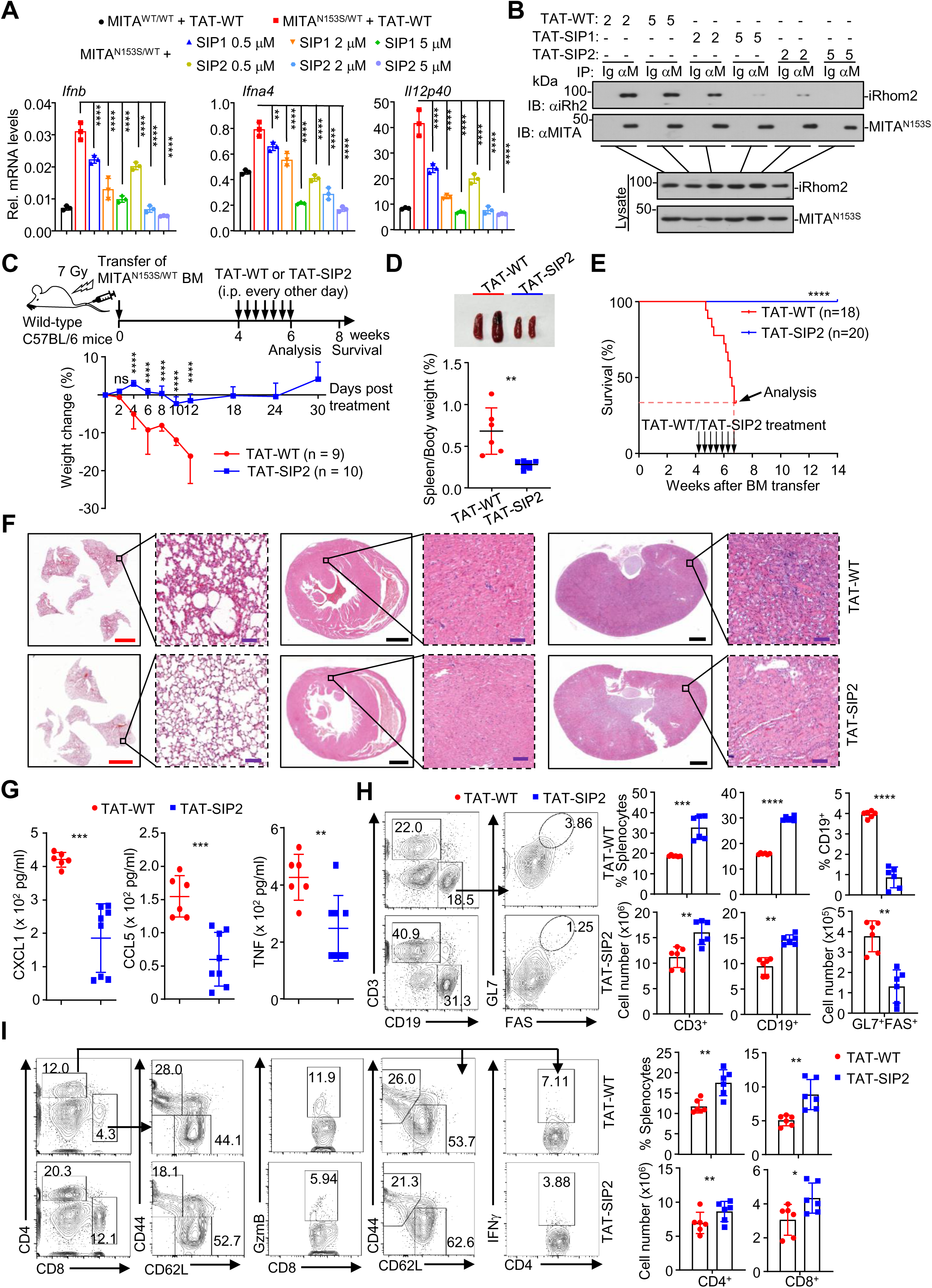
The SAVI inhibitory peptide abrogates the autoimmune SAVI phenotypes in MITA^N153S/WT^ chimeric mice. (A) RT-qPCR analysis of *Ifnb, Ifna4,* and *Il12p40* mRNA levels in MITA^WT/WT^ or MITA^N153S/WT^ MLFs treated with TAT-WT or TAT-SIP1/2 (0.5μM, 2 μM or 5 μM) for 0-24 h. (B) Immunoprecipitation (with control IgG or an anti-MITA antibody) and immunoblot analysis (with anti-iRhom2 and anti-MITA antibodies) of MITA^N153S/WT^ MLFs treated with TAT-WT or TAT-SIP1/2 (2 μM or 5 μM) for 0-24 h. (C) A scheme of TAT-SIP2 treatment (upper) and weight changes (lower) of MITA^N153S/WT^ chimeric mice intraperitoneally injected with TAT-WT (n=9) or TAT-SIP2 (n=10) (20 mg/kg body weight) at the 5th week after bone marrow cell transfer for two weeks. (D) Representative images of spleens (upper) and spleen weight/body weight ratios (lower) of MITA^N153S/WT^ chimeric mice intraperitoneally injected with TAT-WT (n=6) or TAT-SIP2 (n=7) every other day for two weeks starting from the 5th week after bone marrow cell transfer. (E) Survival (Kaplan–Meier curves) of MITA^N153S/WT^ chimeric mice intraperitoneally injected with TAT-WT (n=18) or TAT-SIP2 (n=20) as described in (D). (F) Representative images of HE-stained heart, kidney, and lung sections from MITA^N153S/WT^ chimeric mice treated with TAT-WT (n=3) or TAT-SIP2 (n=3) as described in (D). (G) ELISA of CXCL1, CCL5 and TNF in the sera of MITA^N153S/WT^ chimeric mice treated with TAT-WT (n=6) or TAT-SIP2 (n=8) as described in (D). (H-I) Flow cytometric analysis of the splenocytes from MITA^N153S/WT^ chimeric mice treated with TAT-WT (n=6) or TAT-SIP2 (n=6) as described in (D). \**P* <0.05, \*\**P*<0.01, \*\*\**P*<0.001, \*\*\*\**P*<0.0001, ns, not significant; (one-way ANOVA, the two-tailed Student’s t-test, or the log-rank test). Scale bars represent 1 mm (black), 100 μm (purple), or 3 mm (red) in (F). The graphs show mean ± SD. The data are combined results from two independent experiments (E) or representative of two independent experiments (A-C, D, F-I).

To examine whether SIP binds to iRhom2 to impede its association with MITA GOFs, we synthesized biotinylated SIP2 and performed in vitro pulldown assays. The results suggested that biotin-SIP2 pulled down HA-iRhom2 in HEK293 cells (Figure S7I). In addition, we observed that a portion of Cy5-labeled TAT-SIP2 was colocalized with HA-iRhom2 in HeLa cells and that Cy5-TAT-SIP2 treatment impeded iRhom2-MITA^N153S^ colocalization (Figure S7J). Expectedly, TAT-SIP2 inhibited the Golgi localization of MITA^N153S^ (Figure S8A). In contrast, cGAMP-induced ER-to-Golgi translocation of wild-type MITA or the Golgi localization of MITA^R283S^ was not affected by TAT-SIP2 (Figure S8B-C). Importantly, although treatment with TAT-SIP2 impaired the expression of *Ifnb*, *Ifna4* and *Il12p40* in MITA^N153S/WT^ MLFs (Figure 7A and Figure S8D), such a treatment did not affect HSV-1-induced upregulation of *Ifnb*, *Cxcl10*, *Isg56* and *Il-6* in MITA^N153S/WT^ MLFs (Figure S8D), indicating that SIP does not interfere with the antiviral activity of MITA. Collectively, these data suggest that the designed SIPs selectively inhibit the Golgi localization and activation of MITA hinge-region GOFs by impeding their association with iRhom2.

### 2.9. SIP treatment alleviates the autoimmune SAVI phenotypes in mice

We next examined whether treatment with SIP inhibited the progression of SAVI in mice. As shown in Figure 7C and D, intraperitoneal injection of TAT-SIP2 but not TAT-WT (20 mg/kg body weight) maintained the body weight of and substantially prevented splenomegaly in MITA^N153S/WT^→WT chimeric mice. Strikingly, all the MITA^N153S/WT^→WT chimeric mice that received TAT-SIP2 survived till the 14^th^ week after bone marrow transfer, whereas the same mice that received TAT-WT began to die at 4.5 weeks after bone marrow transfer and more than 60% died at the end of treatment (Figure 7E). Expectedly, the systemic inflammation as determined by the infiltration of immune cells and the expression of proinflammatory cytokines in various organs and the production of CXCL1, CCL5 and TNF in the blood of MITA^N153S/WT^→WT chimeric mice was significantly compromised by TAT-SIP2 treatment (Figure 7F-G and Figure S8E). Consistently, treatment with TAT-SIP2 substantially restored the immune cell development in various lymphoid organs including the bone marrow, thymus, and spleen of the MITA^N153S/WT^→WT chimeric mice (Figure 7H-I and Figure S8F-H). Taken together, these data suggest that targeting the MITA GOFs-iRhom2 interaction by SIP alleviates autoimmune SAVI phenotypes, which provides a potential therapeutic strategy for SAVI patients.

## 3. Discussion

SAVI is a rare autoimmune disease caused by MITA GOFs and the mutated sites are clustered in the hinge region [V147L/M (Abid et al., 2020; Dai et al., 2020), N154S and V155M (Jeremiah et al., 2014; Liu et al., 2014), F153V and G158A(Lin et al., 2021)] and the LBD [C206Y/G (Melki et al., 2017), G207E (Keskitalo et al., 2019), F269S (Valeri et al., 2024), F279L (Seo et al., 2017), R281Q/W (Melki et al., 2017; Volpi et al., 2019), and R284S (Konno et al., 2018; Melki et al., 2017; Saldanha et al., 2018)]. It is acknowledged that mutations in the LBD (F269S, F279L, R281Q/W, and R284S) disrupt the autoinhibition of MITA and thereby lead to its spontaneous activation (Ergun et al., 2019; Liu et al., 2023), whereas how mutations in the hinge region result in the activation of MITA remains unclear. Here, we have provided biochemical and genetic evidence that MITA hinge-region GOFs constitutively interact with iRhom2 for ER-to-Golgi translocation and activation in a manner dependent on Lys150. Mutation at Lys150 of MITA hinge-region GOFs suppressed the GOF-caused destabilization of the inactive state, impaired their associations with iRhom2, inhibited their ER-to-Golgi translocation, and attenuated their capability of activating the expression of type I IFNs and proinflammatory cytokines. In addition, the SAVI autoimmune phenotypes including systemic inflammation and dysregulated development of immune cells were abolished in the MITA^NS/NS^ mice or the *iRhom2*^-/-^MITA^N153S/WT^→WT chimeric mice compared to the respective controls. Meanwhile our structural analysis established that the N154S mutation resulted in an outward shift of the hinge region of MITA, which was further destabilized by K150 and iRhom2. These findings together reveal a previously uncharacterized mechanism of the activation of the MITA hinge-region GOFs.

It has been reported that multiple E3 ligases catalyzed post-translational modifications (PTMs) on Lys150 of MITA after HSV-1 infection or dsDNA challenge. For example, HERC5/6 and TRIM56 mediate MITA Lys150 ISGylation and K63-linked ubiquitination, respectively (Qin et al., 2024; Tsuchida et al., 2010), while RNF26 promotes MITA stabilization via catalyzing K11-linked ubiquitination at Lysl50, counteracting RNF5-dependent K48-linked ubiquitination (Qin et al., 2014; Zhong et al., 2009). However, we have demonstrated that MITA^K150R^ activates ISRE luciferase to a similar extent as wild-type MITA in reporter assays (Zhong et al., 2009), and found that MITA^K150N^ bound to cGAMP, translocated from ER to Golgi apparatus, and upregulated *Ifnb*, *Il6*, *Isg56* and *Cxcl10* after cGAMP treatment as similarly as wild-type MITA, which indicates that Lys150 is dispensable for the cGAMP-induced activation of MITA. It should be noted that the above E3s that catalyze PTMs of MITA also modify other innate immune components such as cGAS [HERC5-mediated ISGylation (Chu et al., 2024); TRIM56-mediated ubiquitination (Seo et al., 2018)] and IRF3 [HERC5-mediated ISGylation (Shi et al., 2010); RNF26-mediated ubiquitination (Qin et al., 2014)]. This implies that their regulatory effects likely extend beyond MITA alone, collectively enhancing cellular antiviral responses through PTMs of multi-substrate proteins. In addition, other lysine residues also undergo the same PTMs by distinct E3s. For example, RNF115 mediates K63-linked ubiquitination at K20, K224, and K289 (Zhang et al., 2024, 2020), whereas TRIM29 catalyzes K48-linked polyubiquitination at K370 (human MITA) and K288/K337 (mouse MITA) (Li et al., 2018a; Xing et al., 2017).

Multiple residues (K224, K236, K289, K338, K347, K370) of MITA are ISGylated, with ISGylation at K289 being critical for MITA function (Lin et al., 2023). Such multi-site modifications may confer redundancy or compensatory mechanisms to ensure MITA activation and downstream signaling even when individual residues are perturbed. The functional interplay among these sites, including potential crosstalk with K150, remains an important area for future investigation.

We have previously shown that iRhom2 interacts with MITA and mediates the ER-to-Golgi translocation and activation of MITA after cGAMP treatment or HSV-1 infection (Luo et al., 2016). In the present study, we found that iRhom2 also interacted with MITA^NS^ after HSV-1 infection and that knockdown of iRhom2 inhibited ER-to-Golgi translocation of MITA^NS^ and impaired expression of type I IFNs and proinflammatory cytokines in MITA^NS/NS^ MLFs after cGAMP treatment. In addition, cGAMP induced the expression of downstream type I IFNs and proinflammatory cytokines in *Mita*^-/-^ cells reconstituted with MITA^K150N^ or MITA^NS^ to similar levels as in *Mita*^-/-^ cells reconstituted with wild-type MITA, suggesting that Lys150 of MITA is dispensable for cGAMP-induced association with iRhom2 and expression of downstream genes. In this context, we found that cGAMP bound to MITA^NS^ as similarly as it did to wild-type MITA. Collectively, these observations imply that the mechanism of activation of MITA hinge-region GOFs, which is dependent on both iRhom2 and Lys150 of MITA, is distinct from that of cGAMP-induced activation of MITA, which is dependent on iRhom2 but independent of Lys150 of MITA.

Our recent structural analyses suggest that two MITA molecules form a dimer through the hinge region and four or more MITA dimers form a side-by-side and head-to-head packing of MITA under homeostatic conditions, thereby holding MITA on two folded ER membranes in an autoinhibitory conformation (Liu et al., 2023). Binding to cGAMP induces the hinge region of each MITA LBD to undergo inward rigid body rotation, which traps the ligand using four-stranded β sheets to generate what is known as the closed conformation (Gao et al., 2013; Zhang et al., 2019, 2013). Mutations in the LBD domain (F269, F279, R281, and R284) disrupt the autoinhibition stability of MITA multimers, leading to its spontaneous activation. This process, however, does not depend on K150 in the hinge region. It has been speculated that mutations at the hinge region might cause a similar rotation as the cGAMP binding for the activation of MITA, which lacks direct structural evidence due to the difficulties in obtaining stable MITA GOF proteins. The results from our structural analysis suggested that two MITA^NS^ molecules formed a similar crossover at the hinge region to wild-type MITA, suggesting that K150 promotes the destabilization of the hinge-region crossover and the activation of MITA^N154S^ in a manner dependent on iRhom2. In addition, compared to wild-type apo-MITA, N152 and E148 in the hinge region formed new interactions with other residues and the connector loop of the hinge region shifted outward in apo-MITA^NS^. Together with the observations that MITA hinge-region GOFs interacted with iRhom2, which was impaired by further mutation at Lys150, these findings indicated that the gain-of-function mutations at the hinge-region and the Lys150 of MITA might provide a docking conformation for iRhom2 that promoted the destabilization and activation of MITA hinge-region GOFs as well as the subsequent ER-to-Golgi trafficking.

There is no effective and specific treatment for SAVI in clinical practice. Although treatment with JAK inhibitors (ruxolitinib, baricitinib, and tofacitinib) results in partial alleviation in ∼50% of SAVI patients, the other half respond poorly to such treatment, indicating the IFN-JAK-STAT pathways partially contribute to the progression of SAVI. In this context, it has been observed that knockout of IRF3 or IFNAR1 does not rescue the SAVI phenotypes in MITA^N153S/WT^ mice (Luksch et al., 2019; Warner et al., 2017). In addition, lung fibrosis and infections have been observed in both responsive and nonresponsive patients after treatment with JAK inhibitors (Balci et al., 2020; Frémond et al., 2021, 2016; Sanchez et al., 2018; Volpi et al., 2019; Wang et al., 2021). Therefore, there is an urgent need to develop selective and precise therapeutic strategies for SAVI. Inspired by the above notion, we synthesized SIP peptides spanning the mutated site and Lys150 in the hinge region of MITA to selectively inhibit the activation of MITA hinge-region GOFs by impeding their associations with iRhom2. We found that SIP selectively impaired the expression of proinflammatory cytokines by MITA hinge-region GOFs and attenuated the SAVI phenotypes of MITA^N153S/WT^→WT chimeric mice without affecting cGAMP-induced activation of wild-type MITA. In addition, the SIP peptides harboring the N153S (for mouse) or N154S (for human) mutations and the Lys150 bound to iRhom2 in cells and *in vitro* and impaired the association between iRhom2 and MITA^N153S^. Notably, SIP-treated MITA^N153S/WT^ MLFs still responded to HSV-1 infection, indicating that selective disruption of the associations between iRhom2 and MITA hinge-region GOFs tunes down SAVI inflammation without affecting the normal antiviral immune responses. Further investigations are needed to reveal how SIP binds to iRhom2 and whether the SIP-binding pocket of iRhom2 serves as a druggable site for the future treatment of SAVI patients. Taken together, these findings suggest diverse activation mechanisms of cGAMP-bound MITA and MITA GOFs and highlight a potential targeted therapeutic strategy for SAVI.

## 4. Methods and materials

### 4.1. Mice

The MITA^N153S/WT^ mice were generated by GemPharmatech Co., Ltd. and maintained as previously described (Zhang et al., 2024). Wild-type C57BL/6 mice were purchased from GemPharmatech Co., Ltd. MITA^NS/NS^ mice were generated similarly through CRISPR/Cas9-mediated gene editing. In brief, guide RNA (5’-GTTAAATGTTGCCCACGGGC-3’) was synthesized through in vitro transcription and purification. The gRNA was incubated with purified Cas9 protein and injected into fertilized eggs (at the one-cell stage) together with the donor vector containing the target mutations in the *Mita* gene. The injected fertilized eggs were cultured to the two-cell stage and subsequently transplanted into pseudopregnant mice. The targeted genomes of F0 mice were amplified via PCR and sequenced and the positive F0 mice were crossed with wild-type C57BL/6 mice to obtain F1 MITA^NS/WT^ mice which were further crossed to obtain the homozygous MITA^NS/NS^ mice. The MITA^NS/NS^ mice were crossed for maintenance. The *Mita*^-/-^ and *iRhom2*^-/-^ mice were described previously and kindly provided by Dr. Hong-Bing Shu (Wuhan University) (Li et al., 2012; Luo et al., 2016). For the generation of *iRhom2*^-/-^MITA^N153S/WT^ mice, wild-type C57BL/6 mice whose ovaries were removed and replaced with the ovaries of MITA^N153S/WT^ mice were crossed with *iRhom2*^-/-^mice to generate the *iRhom2*^+/-^MITA^N153S/WT^ mice. The ovaries of the *iRhom2*^+/-^MITA^N153S/WT^ mice were transplanted to wild-type C57BL/6 followed by crossing with *iRhom2*^-/-^ mice to generate the *iRhom2*^-/-^MITA^N153S/WT^ mice. The *iRhom2*^-/-^MITA^N153S/WT^ mice were maintained by crossing the *iRhom2*^-/-^ mice with the wild-type C57BL/6 mice transplanted with the ovaries of the *iRhom2*^-/-^MITA^N153S/WT^ mice. Age- and sex-matched wild-type, MITA^N153S/WT^, MITA^NS/NS^, *iRhom2^-/-^*MITA^N153S/WT^ mice were used for all the described experiments. All the mice were housed in the specific pathogen-free animal facility at Wuhan University, and all the animal experiments were carried out following protocols approved by the Institutional Animal Care and Use Committee of Wuhan University (Approval No. 24020A). The primers for genotyping were as follows: MITA^N153S^: 5′-AAGAGGTGGGTGGTGTGGGAAG-3′ (forward) (for PCR and sequencing) and 5′-CCTACCTGGTAAGATCAACCGCAAG-3′ (reverse). Wild-type allele of *iRhom2*: 5′-CTCTGAACTTGGGAGTTCTGTTGCC-3′ (forward) and 5′-TTCTAGCAAGGTCTGGGGATACAGC-3′ (reverse); disrupted allele of *iRhom2*: 5′-GAGATGGCGCAACGCAATTAATG-3′ (forward) and 5′-CCATGGTGCTCGTGTTTATGGAAGG-3′ (reverse).

### 4.2. Protein expression and purification

Protein expression and purification of human MITA^NS^ were performed as previously described (Liu et al., 2023). Briefly, the coding sequence for MITA^NS^ was cloned into a modified pEZT-BM mammalian vector featuring a human rhinovirus 3C protease cleavage site and a C-terminal T6SS-secreted immunity protein 3 (Tsi3) tag from *Pseudomonas aeruginosa* (Lu et al., 2014). Transient expression was performed in HEK293F cells cultured in SMM 293-TI Expression Medium (Sino Biological), with enhancement via 3 mM sodium butyrate supplementation 12 hours post-transfection. Cells were harvested after 48 hours. For apo-MITA^NS^ purification, cell pellets were resuspended in buffer A (25 mM Tris pH 8.0, 150 mM NaCl, 1 mM AEBSF) and lysed with 1.5% DDM/CHS (5:1 ratio) during 4 hours of stirring. After removing insoluble debris by centrifugation at 100,000 × g for 30 minutes, the supernatant was loaded onto Tse3-conjugated resin pre-equilibrated with buffer B (20 mM Tris pH 8.0, 150 mM NaCl, 1 mM CaCl₂, 0.02% DDM, 0.004% CHS). Following washes with buffer B, bound protein was treated with 3C protease for on-column cleavage at 4°C for 12 hours. Eluted apo-MITA^NS^ was further purified by size-exclusion chromatography on a Superdex 200 Increase 10/300 column (Cytiva) in buffer C (20 mM HEPES pH 7.5, 150 mM NaCl, 0.02% DDM, 0.004% CHS, 1 mM DTT). Peak fractions were pooled and concentrated to 4 mg/ml for cryo-EM analysis. MITA^NS^/cGAMP complex purification employed identical initial steps except that lysis utilized 1.5% DDM, and subsequent buffers contained 0.1% DDM. Following size-exclusion chromatography, detergent was exchanged using amphipol A8-35 (Anatrace) in the presence of 1 µM cGAMP. The detergent-free complex was concentrated to 0.8 mg/ml for storage at -80°C.

### 4.3. Cryo-EM sample preparation and data collection

Cryo-EM sample preparation and data collection followed standard procedures. For apo-MITA^NS^ and the cGAMP-bound MITA^NS^ filament, 3.5 μL of sample (4 mg/mL or 0.8 mg/mL, respectively) was applied to glow-discharged Quantifoil R2/1 holey carbon grids. Grids were blotted for 3 seconds at 100% humidity and plunge-frozen in liquid ethane using a Vitrobot Mark IV (Thermo Fisher). Data collection was performed on a 300 kV Titan Krios transmission electron microscope (Thermo Fisher) equipped with a GIF-Quantum energy filter (Gatan; 20 eV slit width) and a Gatan K2 direct electron detector operating in super-resolution mode. Images were recorded using Serial-EM software at pixel sizes of 0.365 Å (apo-MITA^NS^) and 0.42 Å (cGAMP-bound MITA^NS^ filament), with a dose rate of 15 e⁻/pixel/s and a total exposure of ∼60 e⁻/Å² fractionated into 36 movie frames per micrograph. Defocus ranges were approximately -1.2 to -3.0 μm for apo-MITA^NS^ and -1.0 to -2.0 μm for the cGAMP-bound MITA^NS^ filament.

### 4.4. Image processing

Image processing commenced with drift correction of all movie stacks using MotionCor2 (MotionCor2: anisotropic correction of beam-induced motion for improved cryo-electron microscopy - PubMed, n.d.), generating 2 × binned micrographs. Initial contrast transfer function (CTF) parameters for each micrograph were estimated with CTFFIND4.1(Rohou and Grigorieff, 2015), and micrographs exhibiting a resolution limit worse than 6 Å were discarded. Approximately 2000 particles were manually picked from selected micrographs to generate templates for automated particle picking against the entire dataset. Subsequent processing utilized RELION (Zivanov et al., 2018) and cryoSPARC (Punjani et al., 2017).

For the apo-MITA^NS^ dataset, 2,049,672 particles were auto-picked from 17,820 micrographs. Extracted particles underwent three rounds of reference-free 2D classification in cryoSPARC. Particles from well-defined classes (842,189 particles) were selected, and an initial model was generated using cryoSPARC ab initio reconstruction. This model served as the reference for RELION 3D classification. After two rounds of 3D classification without symmetry, a predominant class containing 106,159 particles displaying clear secondary structure features and accurate alignment was identified. These particles underwent a Non-unifrom refinement with a mask and C2 symmetry applied in cryoSPARC, yielding a 3.9 Å resolution map (FSC = 0.143 criterion; Figure S3).

The cGAMP-bound MITA^NS^ filament dataset was processed similarly. From 1,912,110 auto-picked particles, three rounds of reference-free 2D classification in cryoSPARC removed heterogeneous particles, resulting in a clean dataset of 985,594 particles. Two rounds of 3D classification with C2 symmetry identified a single dominant class containing 125,664 particles. Initial Non-unifrom refinement with C2 symmetry yielded a 3.4 Å map. Applying dose-weighting and CTF refinement, followed by a further round of focus refinement, produced a final reconstruction at 3.23 Å resolution (Figure S4). This map revealed the cGAMP-bound MITA^NS^ polymer containing two complete MITA dimers in the active state.

Local resolution was estimated using ResMap (Kucukelbir et al., 2014).

### 4.5. Model building and refinement

Model building of apo-MITA^NS^ was performed by docking the previously solved apo-MITA dimer structure (PDB ID: 6NT6) into the apo-MITA^NS^ cryo-EM map (PDB ID: 9VXT). Given the high structural identity, the primary modification involved manually building the K150N and N154S mutations into the map using Coot (Emsley et al., 2010). Similarly, the cGAMP/MITA complex structure (PDB ID: 8IK3) was docked into the cGAMP-bound MITA^NS^ filament map (PDB ID: 9VXU), followed by manual building of the K150N and N154S mutations in Coot. Both models underwent real-space refinement against their respective maps using phenix.real_space_refine (Afonine et al., 2018), and final model quality was assessed with MolProbity (Williams et al., 2018) (Table S1). Structure analysis and figure rendering utilized PyMOL (Pymol: an open-source molecular graphics tool – ScienceOpen, n.d.), UCSF Chimera (Pettersen et al., 2004), and UCSF ChimeraX (Meng et al., 2023).

### 4.6. Bone marrow transfer

C57BL/6 wild-type mice (8 weeks old) were lethally irradiated (7 Gy, 3.5 Gy twice) followed by the injection of bone marrow cells (1×10^6^ each mouse) from MITA^N153S/WT^, or *iRhom2^-/-^*MITA^N153S/WT^ mice via the tail vein.

### 4.7. Flow cytometry analysis

Single-cell suspensions prepared from various tissues were resuspended in FACS buffer (PBS, 1% BSA) and blocked with anti-mouse CD16/32 antibodies for 15 min before staining with antibodies against surface and intracellular proteins (BioLegend). To analyze the expression of IFN-γ and GzmB in T cells, cells were stimulated with PMA (Sigma, P8139) and ionomycin (Sigma, I0634) in the presence of GolgiStop (BD Biosciences, 554724) for 5 h followed by surface and intracellular staining with the indicated antibodies. Flow cytometry data were acquired on a FACS Celesta or LSR FortessaX20 flow cytometer (BD Biosciences) and analyzed with FlowJo software 10.0 (TreeStar).

### 4.8. Reagents, antibodies, and constructs

Recombinant mouse GM-CSF (CHAMOT Biotechnology, #CM060-20MP), DMSO (Sigma, #D8418), and cGAMP (InvivoGen, #1441190-66-4), and c[3’-Biotin-16-G (2’,5’) pA (3’,5’) p] (BIOLOG, C196) were purchased from the indicated manufacturers. The following antibodies and control IgG were purchased from the indicated manufacturers: mouse control IgG (Santa Cruz Biotechnology, sc-2025), mouse anti-GOLGA2/GM130 (Proteintech, 11308-1-AP), anti-RHBDF2 (iRhom2) (Abcepta, #AP13588A), anti-FLAG (Sigma, F3165), anti-HA (ABclonal, AE105), anti-GFP (ABclonal, AE012), anti-Tubulin (ABclonal, A12289), and anti-MITA (Cell Signaling Technology, #13647 and ABclonal, A3575). The following staining antibodies for flow cytometric analysis were purchased from the indicated manufacturers: anti-mouse CD19-APC (Biolegend,152409), anti-mouse CD3-FITC (Biolegend,100306), anti-mouse CD3-APC (Biolegend, 100312), anti-mouse FAS-PE (Biolegend, 152608), anti-mouse GL7-Percp (Biolegend, 144610), anti-mouse CD11b-PE (Biolegend,101208), anti-mouse CD11c-Percp (Biolegend,117328), anti-mouse F4/80-BV421 (Biolegend,123137), anti-mouse Ly6G-Apc-Cy7 (Biolegend,127624), anti-mouse CD4-FITC (Biolegend, 100406), anti-mouse CD8-BV510 (Biolegend, 100751), anti-mouse CD44-APC (Biolegend,103012), anti-mouse-CD62L-Percp (Biolegend, 104432), anti-mouse IFNγ-Percp (Biolegend, 505822), anti-mouse GzmB-PE (Biolegend, 372208), anti-mouse CD4-BV421 (Biolegend,100427), anti-mouse Sca-1-APC (Biolegend,108111), anti-mouse lineage-biotin (BD Biosciences, 559971), Streptavidin-APC-Cy7 (BD Biosciences, 554063), and anti-c-Kit-PE (Biolegend, 105807), Cell Counting Kit-8 (CCK-8) (TargetMbl, C0005).

### 4.9. siRNAs

For reporter gene assays, HEK293 cells were transfected with control siRNA (siCon) or siRNAs for iRHOM2 (si *iRHOM2*) together with FLAG-MITA^N153S^ for 36 h followed by luciferase assays. For RT-qPCR assays, MITA^N153S/WT^ MLFs were transfected with t siRNA (si Con) or siRNAs for iRhom2 (si *iRhom2*) for 24 h followed by RT-qPCR analyses. The siRNAs were synthesized by JTSBIO Co., Ltd (Wuhan,China) and the sequences of the siRNAs were as follows: siCon: UUCUCCGAACGUGUCACGUTT; si-h*RHBDF2*(*iRHOM2*) #1: GGAAGAACCCAGCCUACUUTT; si-*hRHBDF2*(*iRHOM2*) #2: GCCUCCAAGGUGAAGCACUTT; si-*iRhom2*: CCAUGCCUGACGAUGUCUUTT.

### 4.10. Reporter gene assays

The experiments were performed as previously described (Gao et al., 2025). HEK293T cells were seeded into 24-well plates at 2 × 10^5^ cells per well. HEK293T cells were transfected with the IFN-β promoter luciferase reporter plasmid (0.15 μg), the TK Renilla luciferase control plasmid (0.05 μg) and wild-type MITA or MITA GOFs (0.15 μg). Twenty-four hours later, luciferase assays were performed with a dual-specific luciferase reporter kit (Promega).

The activity of firefly luciferase was normalized by that of Renilla luciferase to obtain the relative luciferase activity.

### 4.11. Cell culture

HEK293 cells were obtained from the American Type Culture Collection, authenticated by short tandem repeat (STR) profiling, and tested for mycoplasma contamination (Sun et al., 2017). Primary MLFs were isolated from 8 to 10-week-old mice. The lungs were minced and digested in calcium and magnesium-free HBSS buffer supplemented with type I collagenase (10 mg/ml) (Worthington) and DNase I (Sigma‒Aldrich) (20 μg/ml) for 2-3 h at 37 °C with shaking. The cell suspensions were filtered and cultured in DMEM supplemented with 10% (vol/vol) FBS and 1% streptomycin-penicillin. Two days later, adherent fibroblasts were rinsed with PBS and subjected to various analyses or assays.

### 4.12. Coimmunoprecipitation and immunoblotting

HEK293 cells were transfected with HA-EXOC2, HA-SAR1a, HA-iRhom2, HA-SEC24c, HA-SURF4, HA-STEEP, HA-STIM1, HA-TRAPβ or HA-TMED2 along with FLAG-hMITA^N154S^, FLAG-hMITA^NS^, mMITA^N153S^ or mMITA^NS^. Twenty-four hours later, cells were collected and lysed on ice for 10 min with 1 mL of Nonidet P-40 lysis buffer (20 mM Tris HCl, pH 7.4-7.5; 150 mM NaCl; 1 mM EDTA; and 1% Nonidet P-40) containing protease and phosphatase inhibitors (APExBIO, K1015). The cell lysates (800 μl) were incubated with control IgG or anti-FLAG antibodies and protein G agarose (20 μl) and rotated shaking for 2 h. For endogenous immunoprecipitation assays, MITA^N153S/WT^ and MITA^NS^ MLFs were collected and lysed as described above. The cell lysates (800 μl) were incubated with control IgG or an anti-MITA antibody and protein G agarose (20 μl) for overnight rotation shaking at 4℃. The immunoprecipitates were washed three times with 1 ml of pre-lysis buffer and subjected to immunoblot analysis. The remaining lysates (200 μl) were subjected to immunoblot analysis with anti-HA, FLAG, Tubulin, MITA or iRhom2.

### 4.13. Immunofluorescence and confocal microscopy analysis

The experiments were performed as previously described (Zhang et al., 2022). MLFs were cultured on coverslips, fixed in 4% paraformaldehyde for 15 min and washed with PBS three times with the front side facing up. The cells were permeabilized with 0.5% saponin in PBS for 5 min and washed with PBS three times. The cells were blocked in blocking buffer (0.1% saponin and 1% BSA in PBS) for 1 h, stained with primary antibodies in blocking buffer (0.1% saponin and 1% BSA) for overnight followed by PBS wash three times (5 min each). The cells were further stained with the Alexa Fluor 488 or 594-conjugated secondary antibodies for 1 h. Finally, the cells were stained with VECTASHIELD® Antifade Mounting Medium with DAPI (Vector, H-1200), and the coverslips were mounted on slides. Images were acquired with a Zeiss LSM880 fluorescence microscope.

### 4.14. Transfection of cGAMP

The cells were treated with penetration buffer (50 mM HEPES, 100 mM KCl, 85 mM Sucrose, 0.2% BSA, 3 mM MgCl2, 0.1 mM DTT, 1 mM ATP, 0.1 mM GTP) (500 μl) containing cGAMP (1 μg) and digitonin (5 μg) for 30 min. Subsequently, the penetration buffer was removed, and complete medium was added for 4 h followed by various assays.

### 4.15. cGAMP binding ability analysis

The experiments were performed as previously described (Li et al., 2018b). HEK293T cells transfected with FLAG-tagged MITA, MITA^K150N^, MITA^N153S^, MITA^NS^, or MITA^R237A^ were harvested and lysed in Nonidet P-40 lysis buffer (20 mM Tris HCl, pH 7.4-7.5; 150 mM NaCl; 1 mM EDTA; and 1% Nonidet P-40) containing protease and phosphatase inhibitors (APExBIO, K1015) with brief sonication. The cell lysates were cleared at 12,000 g for 10 min, and the supernatants were collected and equally divided into two parts, one of which was incubated with biotin (5 μM), and the other was incubated with biotin-cGAMP (5 μM) (BIOLOG, C196-001) at 4℃ overnight, respectively. After incubation, equivalent streptavidin-conjugated agarose beads were added and rotated for 2 h. The beads were washed three times with the Nonidet P-40 lysis buffer, and the bead-bound proteins were eluted and separated by SDS-PAGE for immunoblot analyses.

### 4.16. Hematoxylin-eosin (HE) staining

Various organs including lungs, hearts and kidneys from the indicated mice were fixed in 4% paraformaldehyde for 4 h followed by successive dehydration in 75% ethanol, 95% ethanol, absolute ethanol and xylene (2 h each step). The organs were immersed in a paraffin tank for 48 h and embedded in paraffin blocks. The paraffin blocks were sectioned (6-8 μm) for H&E staining (Beyotime Biotech) followed by cover slipping. Images were acquired using an Aperio VERSA 8 (Leica) multifunctional scanner.

### 4.17. Lentivirus-mediated gene transfer

The experiments were performed as previously described (Guo et al., 2022; Zhang et al., 2024). HEK293 cells were transfected with phage-MITA^WT^-FLAG, phage-MITA^N153S^-FLAG, phage-MITA^K150N^-FLAG, phage-MITA^NS^-FLAG, phage-MITA^V146L^-FLAG, phage-MITA^V154M^-FLAG or phage-MITA^R283S^along with the packaging plasmid psPAX2 and the envelope plasmid pMD2G with PEI reagent (Poly Sciences, 24765-1). The medium was changed to fresh complete medium (10% FBS, 1% streptomycin-penicillin) at 6 h after transfection. Forty hours later, the supernatants were harvested to infect primary *Mita*^-/-^ or *iRhom2*^-/-^ MLFs, followed by various analyses.

### 4.18. RT-qPCR and ELISA

These experiments were performed as previously described (Zhang et al., 2024; Zeng et al., 2026; Yang et al., 2025). Total RNA was extracted from MLFs (2 × 10^6^) or tissues (0.5 mg of liver or brain tissue) using TRIzol (Life Technologies, 15596026). The tissue was minced and homogenized with an IKA T10 basic homogenizer in 1 ml of TRIzol. Chloroform (200 μL) was added to the homogenate, and mixed by inverting it up and down followed by centrifugation at 12,000 × g for 10 min at 4°C. The upper aqueous phase was transferred to a new 1.5 ml Eppendorf tube, and isopropanol (400 μL) was added. Subsequently, the tube was centrifuged at 12,000 g for 10 min at 4°C to obtain a white pellet at the bottom. The pellet was washed twice with 1 ml of 75% ethanol and saved for reverse transcription with All-in-One cDNA Synthesis SuperMix (Aidlab Biotechnologies) and quantitative PCR analysis. For RNA from cells, cells immersed in 1 ml of TRIzol reagent (Invitrogen) were mixed with 200 μl of chloroform and centrifuged (12,000 × g) for 10 min at 4 °C followed by the processes described above. *Ifnb*, *Isg15*, *Isg56*, *Il6*, *Cxcl10* or *Il12p40* expression levels of cells and *Tnf*, *Ccl2*, *Ccl3* or *Ccl4* expression levels of livers, brains and MLFs were measured with a Bio-Rad CFX Connect system by a fast two-step amplification program with Hieff® qPCR SYBR Green Master Mix (Yeasen Biotechnology, 11201ES08). The expression value obtained for each gene was normalized to that of the gene encoding β-actin. The sequences of the PCR primers used were listed in Table S2. The concentrations of CXCL1, IL12p70, IL-6, CCL5, and TNF in serum or in the supernatants were measured with LEGENDplex™ Mul-Analyte Flow Assay KitMouse Anti-Virus Response Panel (13-plex) with V-Bottom Platefrom (Cat. No. 740622, BioLenged) according to the manufacturer’s instructions.

### 4.19. TAT-SIP treatment

TAT-SIP1/2, TAT-WT/NS and TAT were synthesized by GenScript Inc. with over 95% purity. All the above peptides are acetylated at the N-terminus and amidated at the C-terminus. For in vitro studies, the culture medium was removed at 6 hours after transfection. Fresh complete medium containing the peptides (2 μM or 5 μM) was added for 24 h followed by various experiments. For in vivo studies, the MITA^N153S/WT^→WT chimeric mice at the 5th week after bone marrow transfer were intraperitoneally injected with TAT or TAT-SIP2 (20 mg/kg) every other day for 2 weeks followed by various analyses.

### 4.20. Cell viability test

The cytotoxicity of peptides in MLFs cells was determined by CCK-8 assay. Cells were seeded into 96-well plates for twenty hours, and the peptides were added and incubated at 37°C for 24 h. Subsequently, the cell supernatant was replaced with fresh full DMEM medium, and the CCK-8 solution was added at 37°C for 1-4 h followed by measurement at the absorbance of 450 nm.

### 4.21. Biotin-SIP2 binding analysis

HEK293 cells transfected with HA-iRhom2 were harvested and lysed in Nonidet P-40 lysis buffer. The cell lysates were cleared at 12,000 g for 10 min, and the supernatants were collected and equally divided into two parts, one of which was incubated with biotin (5 μM), and the other was incubated with biotin-SIP2 (5 μM) at 4℃ overnight. After incubation, equivalent streptavidin-conjugated agarose beads were added and rotated for 2 h. The beads were washed three times with the Nonidet P-40 lysis buffer, and the bead-bound proteins were eluted and separated by SDS-PAGE for immunoblot analyses.

### 4.22. Statistical analysis

Differences between the experimental and the control groups were tested using Log-Rank analysis, one-way ANOVA, or Student’s *t*-test. The number of repeats for each experiment is also indicated in the respective Figure legends. The tests used for statistical analyses are described in the legends of each concerned Figure and have been performed using GraphPad Prism 8.3.0 or R 4.1.1. Symbols for significance: ns, non-significant; \**P* <0.05, \*\**P*<0.01, \*\*\**P*<0.001, \*\*\*\**P*<0.0001. The values were expressed as mean ± SD unless otherwise indicated.

## Data availability

The datasets and reagents generated during and/or analyzed during the current study are available from the corresponding author on reasonable request.

## Acknowledgments

We thank Dr. Hong-Bing Shu (Wuhan University) for the reagents and members of the Zhong laboratory and the core facilities of Medical Research Institute for technical help. This study was supported by grants from the National Key Research and Development Program of China (Grant Nos. 2023YFC2306100, 2024YFA1803103 and 2024ZD0524901), the Natural Science Foundation of China (Grant Nos. 82425027, 32470960, 82402062, 324B2028, 32270951, and 823B1006), the Fundamental Research Funds for the Central Universities (Grant No. 2042022dx0003 and 2042025kf0044), the Natural Science Foundation of Wuhan (2024040701010031), the Natural Science Foundation of Hubei (2025AFA026), the Postdoctoral Fellowship Program of CPSF (Grant No. GZC20231975), the Shanxi Provincial Science Fund for Distinguished Young Scholars program (202103021221001 and 202303021223007), and the Doctoral Research Startup Project at Shanxi Agricultural University (SXBYKY2023007).

## Author contributions

B.Z., G.S. and D.L. conceived and supervised the study; F.X.L. and Z.D.Z. performed the experiments and analyzed the data; S.L. and J.Y. performed the Cryo-EM analysis; D.L. helped with the structural analysis; B.K.X. and X.S. helped with the mouse experiments; B.Z., G.S., D.L., F.X.L. and Z.D.Z. wrote the paper. All the authors discussed the results and presented viewpoints.

---

**Figure S1.**
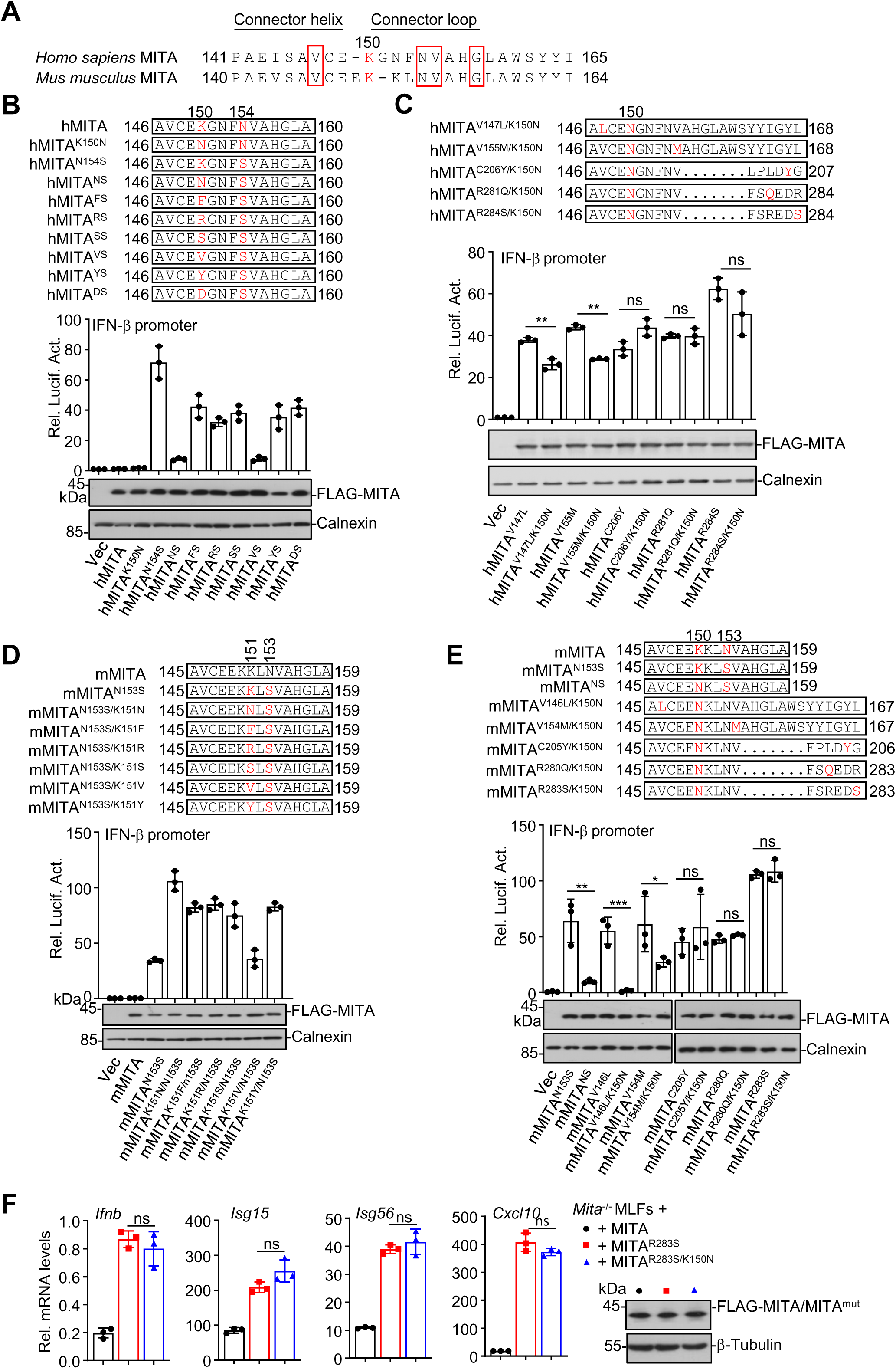
Mutation of K150 impairs the activity of MITA hinge-region GOFs. (A) Sequence alignment of the hinge region of hMITA and mMITA. The amino acid residues in the rectangles are frequently mutated sites for SAVI. (B-E) Luciferase reporter assays of the IFN-β promoter activity and immunoblot assays of the indicated proteins in HEK293 cells transfected with the indicated plasmids, the IFN-β promoter luciferase (0.15 μg) and the internal control TK Renilla luciferase constructs (0.05 μg) for 24 h. (F) RT-qPCR analysis of *Ifnb*, *Isg15*, *Isg56* and *Cxcl10* mRNA levels (left graphs) and immunoblot analysis of the protein levels of MITA, MITA^R283S^ or MITA^R283S/K150N^ (right panels) of *Mita*^-/-^ MLFs reconstituted with MITA, MITA^R283S^ or MITA^R283S/K150N^. \**P* <0.05, \*\**P*<0.01, \*\*\**P*<0.001; ns, not significant (one-way ANOVA). The graphs show mean ± SD. The data are representative of at least three independent experiments (B-F).

**Figure S2.**
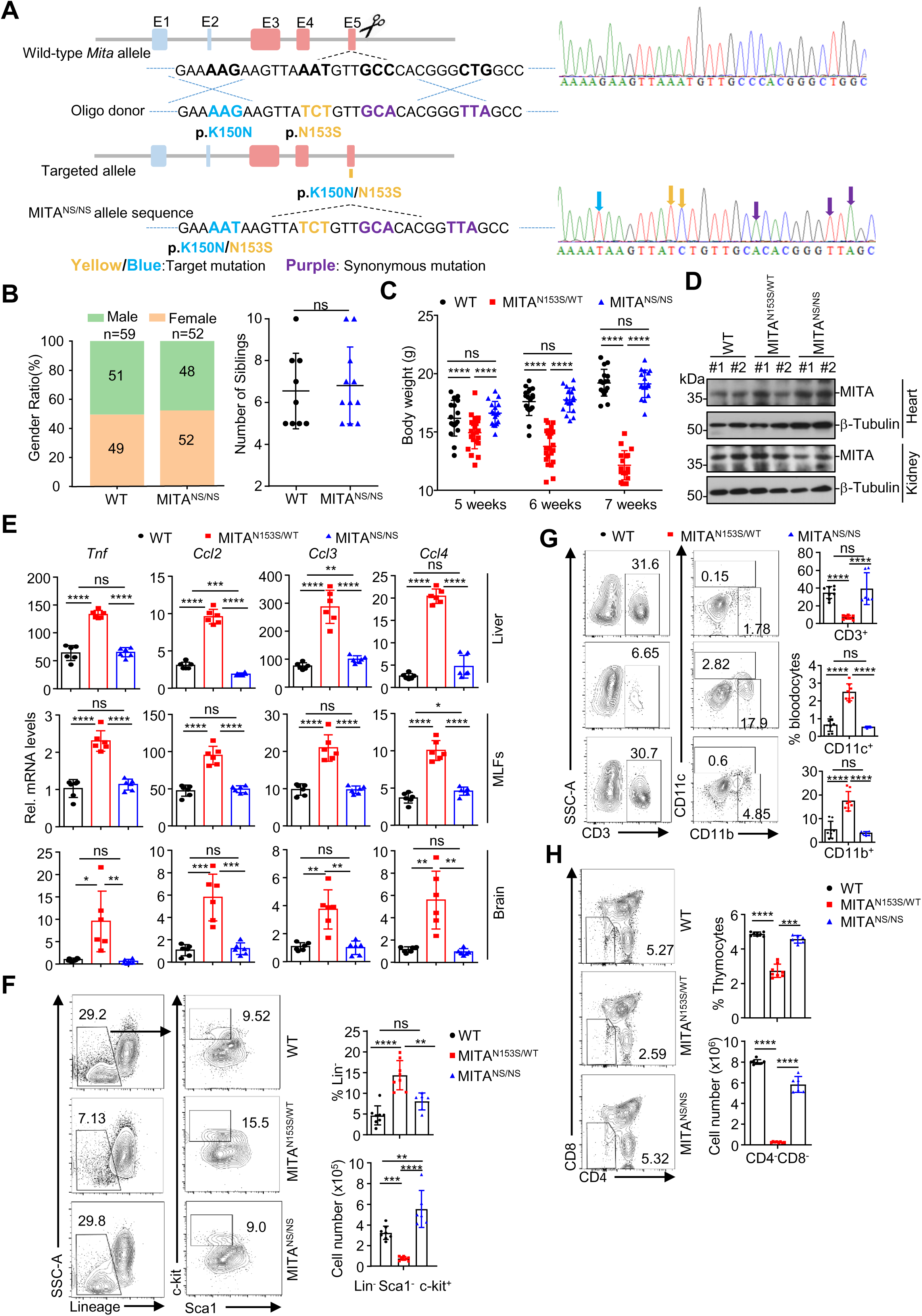
Generation of the MITA^K150N/N153S^ mice. (A) A scheme of CRIPSR/Cas9-mediated genome editing of the *Mita* gene locus. (B) The gender ratio (left) and the number of siblings (right) of WT (n=59) and MITA^NS/NS^ (n=52) mice. (C) Body weights of 5/6/7-week-old WT (n=17), MITA^N153S/WT^ (n=17) and MITA^NS/NS^ (n=17) mice. (D) Immunoblot analysis (with anti-MITA or anti-Tubulin) in hearts and kidneys from WT, MITA^N153S/WT^ and MITA^NS/NS^ mice. (E) RT-qPCR analysis of *Tnf*, *Ccl2*, *Ccl3* and *Ccl4* mRNA in the liver, brain and MLFs of 5-week-old WT(n=6), MITA^N153S/WT^ (n=6) and MITA^NS/NS^ (n=6) mice. (F-H) Flow cytometric analysis of the bone marrow cells (F), peripheral blood cells (G) and thymocytes (H) from 5-week-old WT (n=8), MITA^N153S/WT^ (n=8) and MITA^NS/NS^ (n=6) mice stained with fluorophore-conjugated antibodies against the indicated surface molecules. \**P* <0.05, \*\**P*<0.01, \*\*\**P*<0.001, \*\*\*\**P*<0.0001; ns, not significant (one-way ANOVA or two-tailed student’s t-test). The graphs show mean ± SD. The data are representative of two independent experiments (D-H).

**Figure S3.**
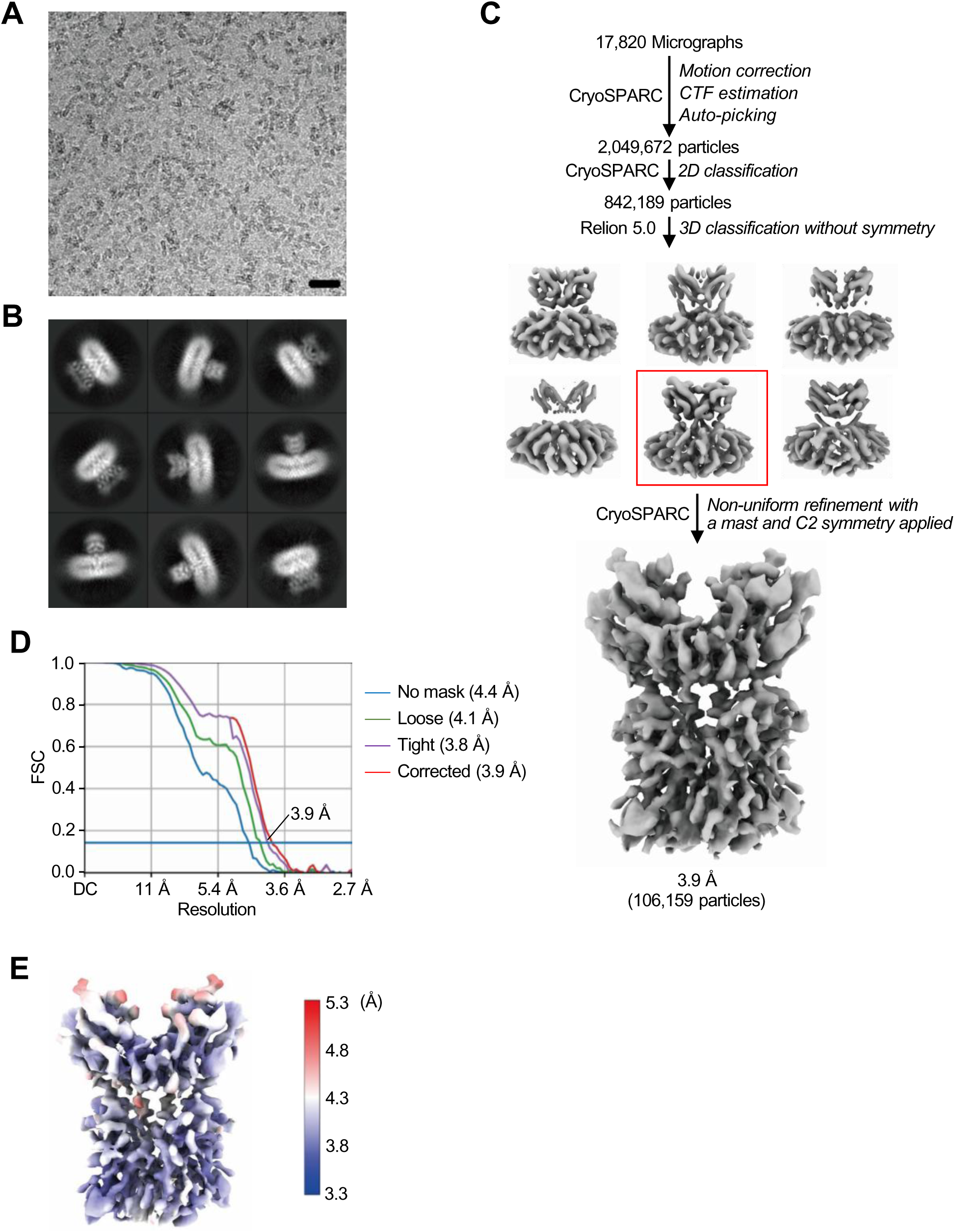
Cryo-EM analysis of apo-MITA^NS^. (A) A representative Cryo-EM micrograph of apo-MITA^NS^. Scale bar, 20 nm. (B) 2D class average images of apo-MITA^NS^. (C) A brief workflow of Cryo-EM image processing and reconstruction. (D) The GSFSC curve for the reconstruction. (E) Local resolution distribution for the density map of apo-MITA^NS^.

**Figure S4.**
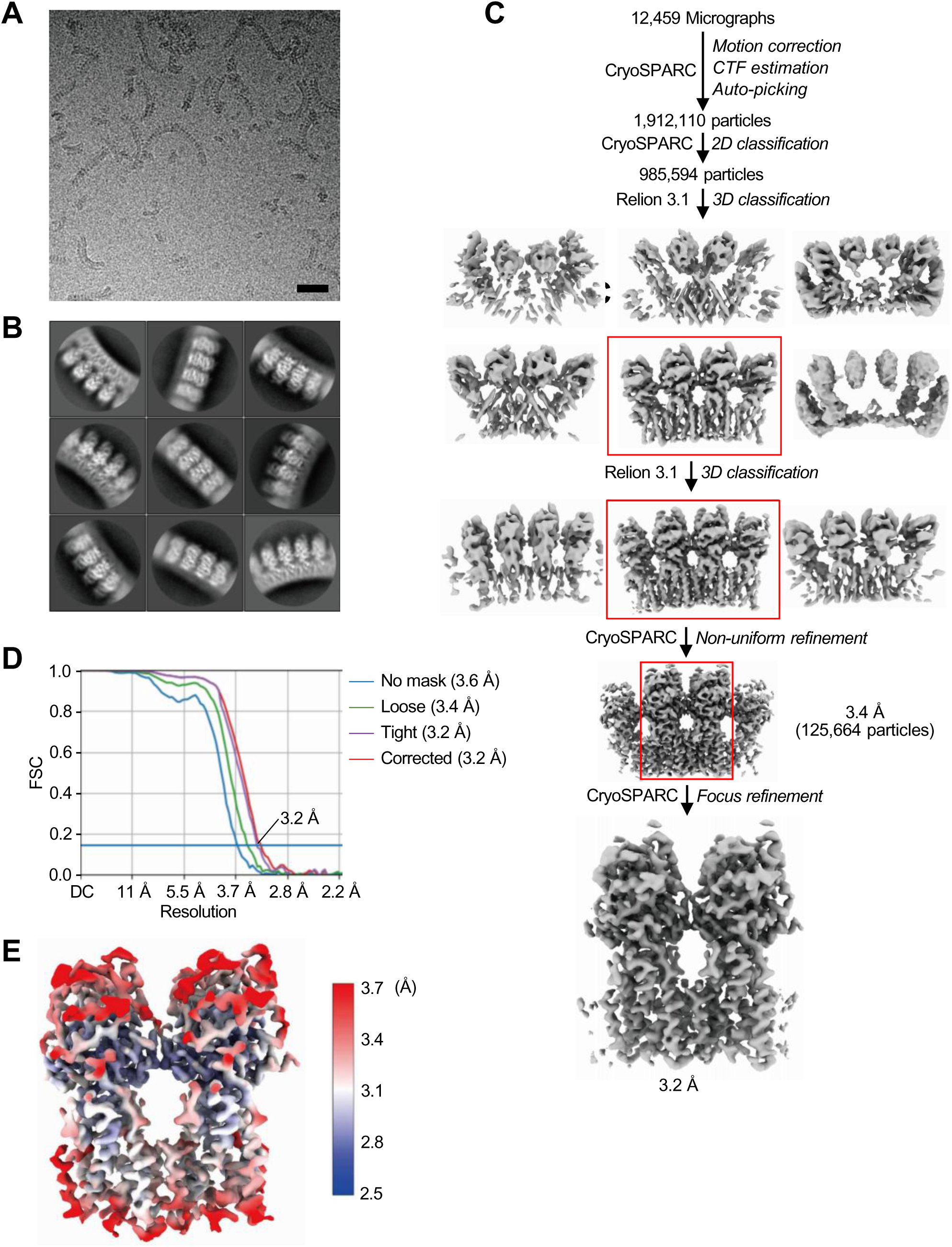
Cryo-EM analysis of cGAMP-bound MITA^NS^. (A) A representative Cryo-EM micrograph of cGAMP-bound MITA^NS^. Scale bar, 20 nm. (B) 2D class average images of cGAMP-bound MITA^NS^. (C) A brief workflow of Cryo-EM image processing and reconstruction. (D) The GSFSC curve for the reconstruction. (E) Local resolution distribution for the density map of cGAMP-bound MITA^NS^.

**Figure S5.**
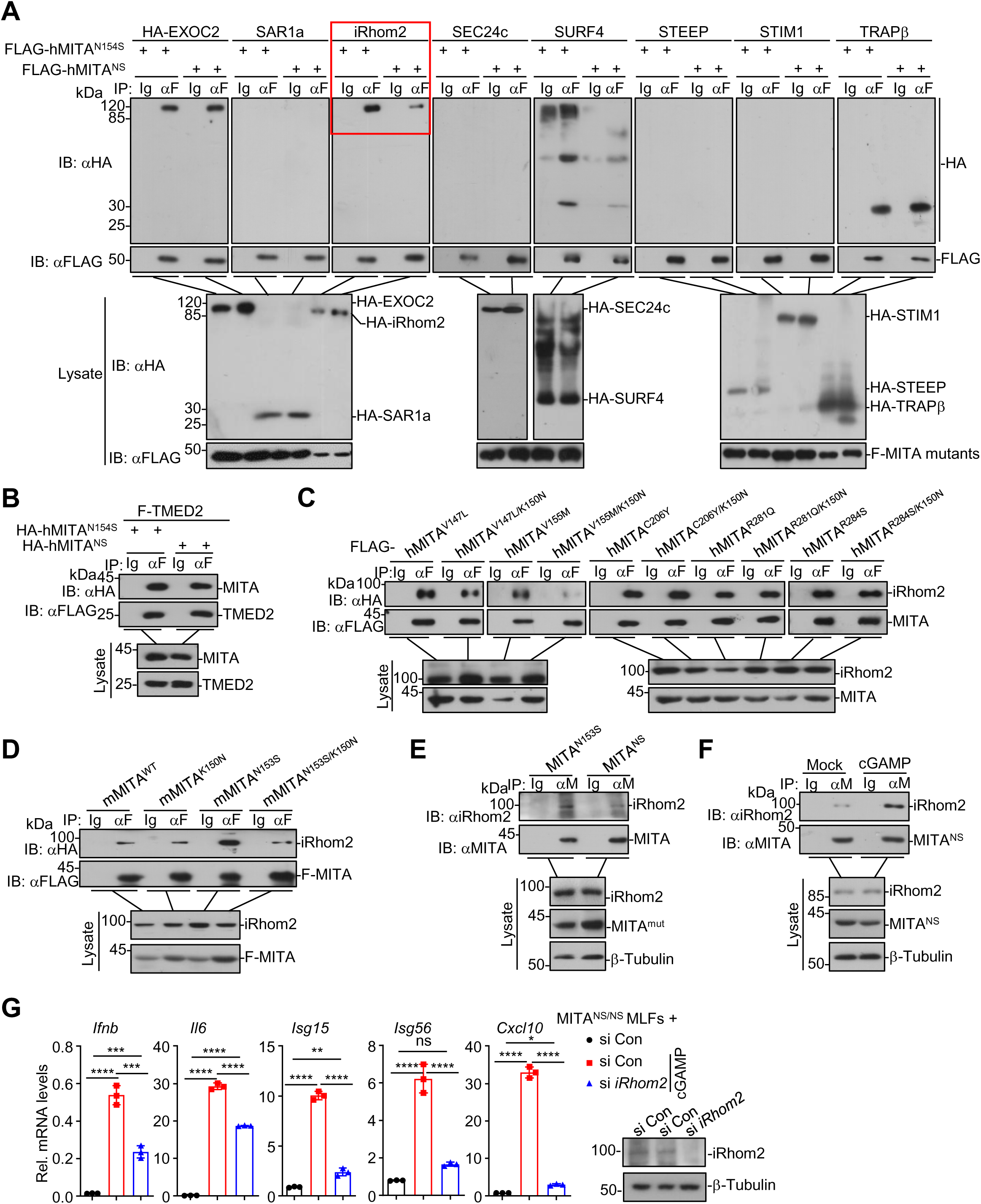
Mutation of K150 in MITA^N153S^ impairs its association with iRhom2. (A-D) Immunoprecipitation (with control IgG or an anti-FLAG antibody) and immunoblot analysis (with anti-FLAG and anti-HA antibodies) of HEK293 cells transfected with the indicated plasmids for 24 h. (E) Immunoprecipitation (with control IgG or an anti-MITA antibody) and immunoblot analysis (with anti-MITA, anti-iRhom2 and anti-Tubulin antibodies) of MITA^N153S/WT^ and MITA^NS/NS^ MLFs. (F) Immunoprecipitation (with control IgG or an anti-MITA antibody) and immunoblot analysis (with an anti-MITA, anti-iRhom2 and anti-Tubulin antibodies) of MITA^NS/NS^ MLFs treated with cGAMP (1 μg/mL) for 0-2 h. (G) RT-qPCR analysis of *Ifnb*, *Il6*, *Isg15*, *Isg56*, and *Cxcl10* mRNA levels (left graphs) and immunoblot analysis of iRhom2 (right panels) of MITA^NS/NS^ MLFs transfect with control siCon or si*iRhom2* followed by treatment with cGAMP (1 μg/mL) for 0-4 h. \**P*<0.05, \*\**P*<0.01, \*\*\**P*<0.001, \*\*\*\**P*<0.0001; ns, not significant (one-way ANOVA). The graphs show mean ± SD. The data are representative of three independent experiments.

**Figure S6.**
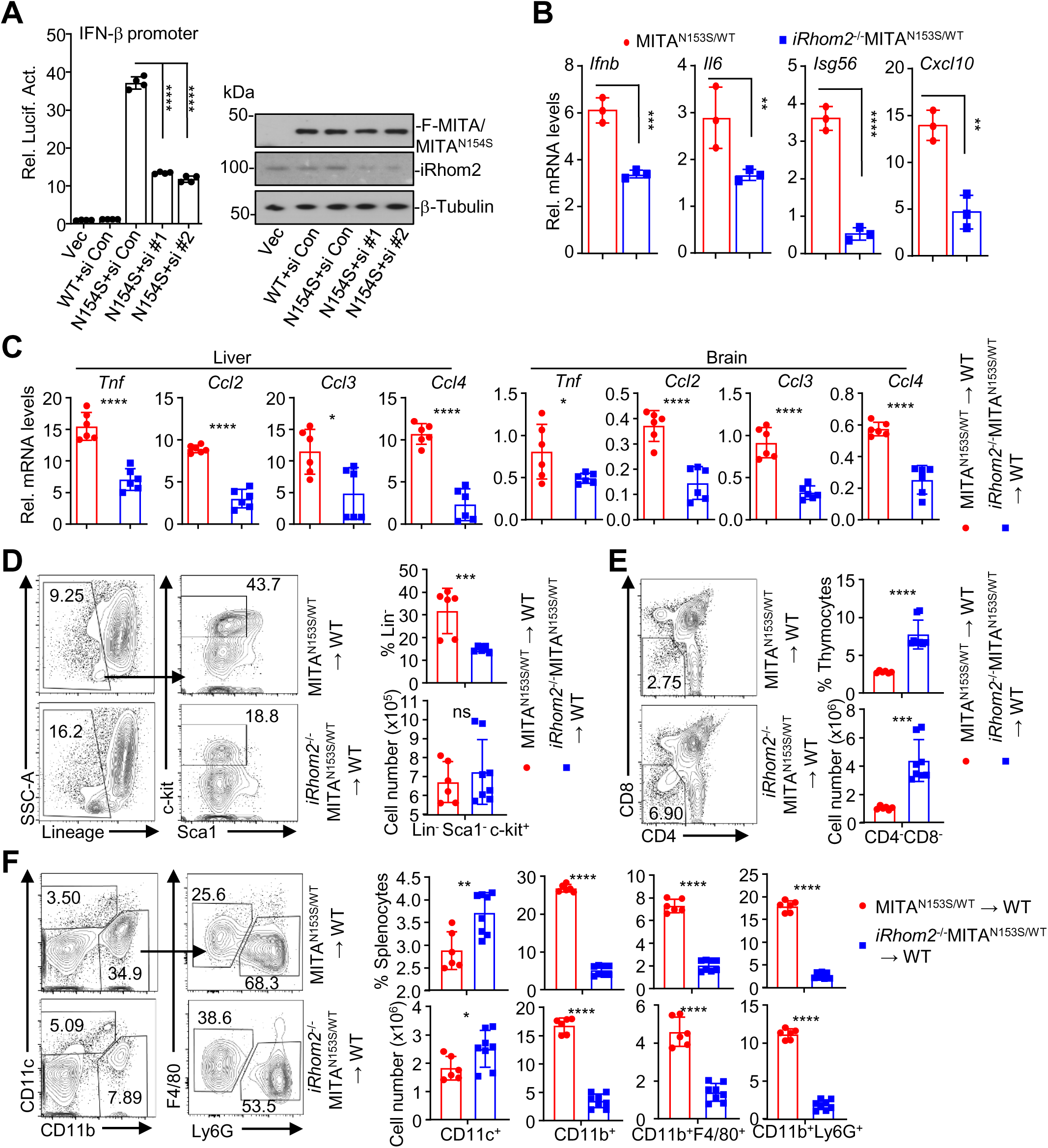
iRhom2 promotes the activation of MITA GOFs. (A). Luciferase reporter assays of IFN-β promoter activity (left graph) and immunoblot analysis of iRhom2, MITA or MITA^N154S^ (right panels) of HEK293 cells transfected with si *iRHOM2* #1, si *iRHOM2* #2 or si Con, together with wild-type MITA or MITA^N154S^ for 36 h. (B) RT-qPCR analysis of *Ifnb*, *Il6*, *Isg56* and *Cxcl10* mRNA levels of MITA^N153S/WT^ or *iRhom2*^-/-^MITA^N153S/WT^ MLFs. (C) RT-qPCR analysis of *Tnf*, *Ccl2*, *Ccl3* and *Ccl4* mRNA levels in the livers and brains of MITA^N153S/WT^→WT (n=6) and *iRhom2*^-/-^ MITA^N153S/WT^→WT (n=6) chimeric mice at the 7th week after bone marrow cell transfer. (D-F) Flow cytometric analysis of bone marrow cells (D), thymocytes (E) and splenocytes (F) from MITA^N153S/WT^→WT (n=6) and *iRhom2*^-/-^MITA^N153S/WT^→WT (n=8) chimeric mice followed by staining with fluorophore-conjugated antibodies against the indicated surface molecules. \**P*<0.05, \*\**P*<0.01, \*\*\**P*<0.001, \*\*\*\**P*<0.0001; ns, not significant (one-way ANOVA or the two-tailed student’s t-test). The graphs show mean ± SD. The data are representatives of three (A) or two (B-F) independent experiments.

**Figure S7.**
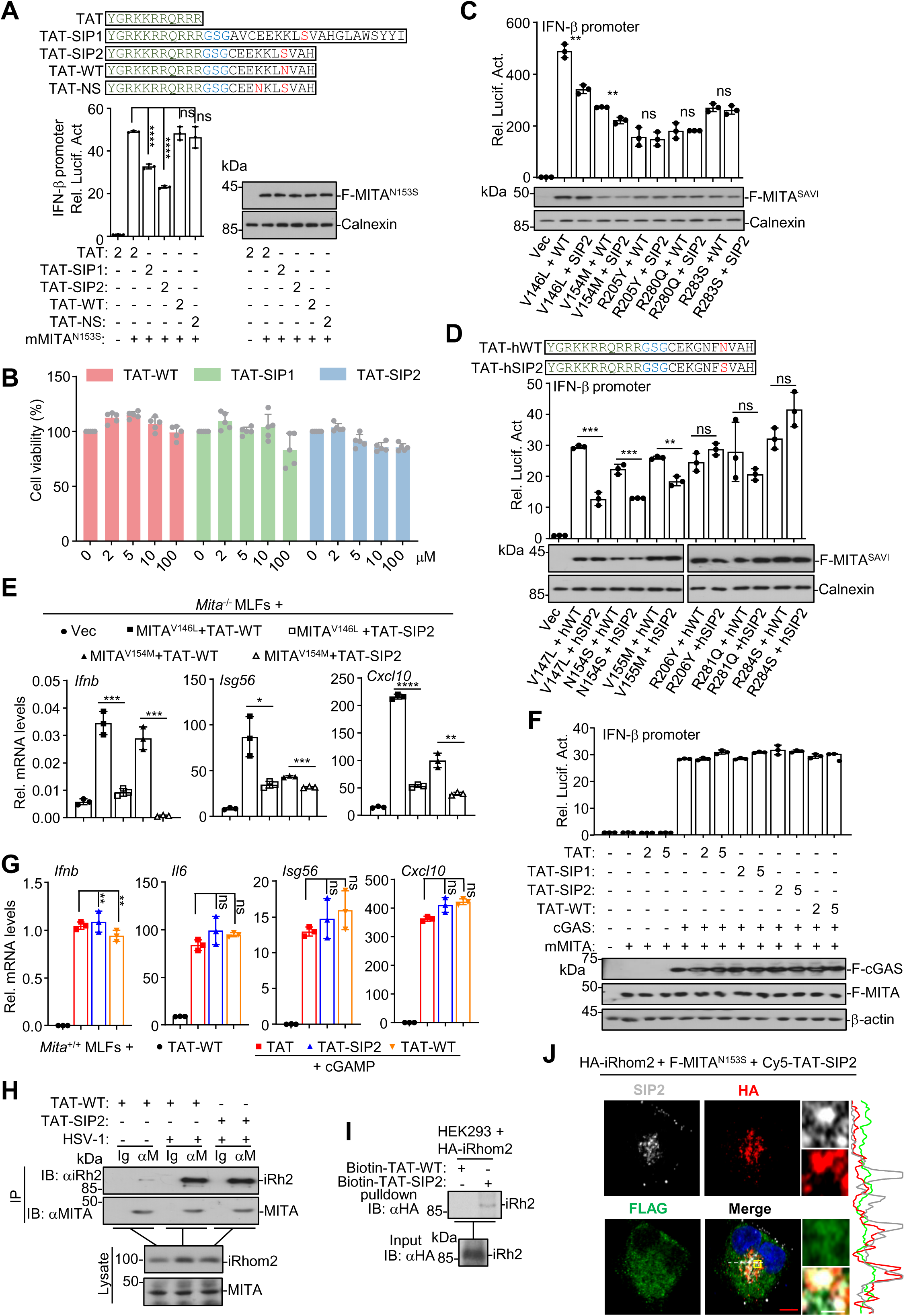
SAVI inhibitory peptides (SIPs) inhibit the activation of MITA GOFs. (A) The sequences of TAT, TAT-SIP1/2 and TAT-WT/NS. Luciferase reporter assays of IFN-β promoter activity of HEK293 cells that were transfected with FLAG-MITA^N153S^ in the presence of TAT, TAT-SIP1/2 or TAT-WT/NS (2 μM) for 24 h. (B) CCK-8 cell viability assays of HEK293 cells treated with the indicated concentrations of TAT-WT, TAT-SIP1 or TAT-SIP2 for 24h. (C) Luciferase reporter assays of IFN-β promoter activity of HEK293 cells transfected with the indicated plasmids followed by treatment with TAT-WT or TAT-SIP2 (2 μM) for 24 h. (D) The sequences of TAT-hWT and TAT-hSIP2 (upper scheme) and luciferase reporter assays of IFN-β promoter activity of HEK293 cells transfected with the indicated plasmids followed by treatment with TAT-hWT or TAT-hSIP2 (2 μM) for 24 h (lower graph). (E) RT-qPCR analysis of *Ifnb*, *Isg56* and *Cxcl10* mRNA levels of *Mita*^-/-^ MLFs reconstituted with MITA^V146L^ or MITA^V154M^ followed by treatment with TAT-WT or TAT-SIP2 (5 μM) for 24 h. (F) Luciferase reporter assays of IFN-β promoter activity of HEK293 cells transfected with cGAS and MITA followed by treatment with TAT-hSIP1/2 or TAT-hWT (2 μM or 5 μM) for 24 h. (G) RT-qPCR analysis of *Ifnb*, *Il6*, *Isg56* and *Cxcl10* mRNA levels of *Mita*^+/+^ MLFs treated with TAT, TAT-SIP2 or TAT-WT (5 μM) followed by transfection with cGAMP (1 μg/mL) for 0-4 h. (H) Immunoprecipitation (with control IgG or an anti-MITA antibody) and immunoblot analysis (with anti-iRhom2 and anti-MITA antibodies) of *Mita*^+/+^ MLFs followed by HSV-1 infection for 0-6 h in the presence or absence of TAT-WT or TAT-SIP1/2 (5 μM). (I) In vitro pulldown assay of iRhom2 that was transfected into HEK293 cells by Biotin-SIP2. (J) Immunofluorescence staining and confocal imaging of HA (red) and FLAG (green) in HeLa cells that were transfected with FLAG-MITA^N153S^ and HA-iRhom2 for 6 h followed by the treatment of Cy5-TAT-SIP2 (5 μM) for 20 h. \**P*<0.05, \*\**P*<0.01, \*\*\**P*<0.001, \*\*\*\**P*<0.0001; ns, not significant (one-way ANOVA). Scale bars represent 1 μm (white) and 5 μm (red) in (J). The graphs show mean ± SD. The data are representative of three (A-D, F) or two (E, G, H-J) independent experiments.

**Figure S8.**
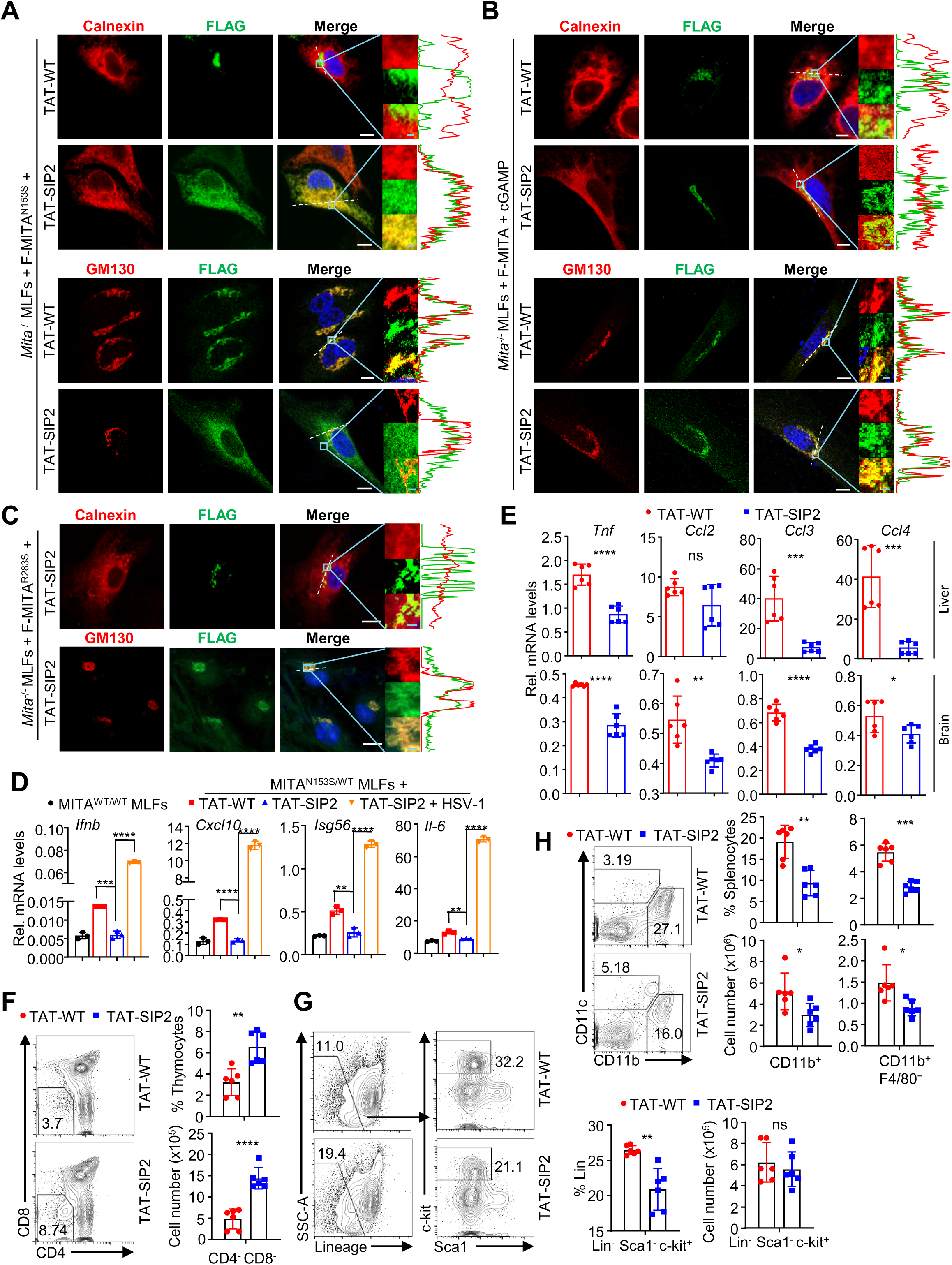
The SAVI inhibitory peptide (SIP) attenuates the autoimmune SAVI phenotypes in MITA^N153S/WT^ chimeric mice. (A) Immunofluorescent staining of Calnexin (an ER marker) (red), GM130 (a Golgi apparatus marker) (red), and FLAG (green) and confocal microscopy analysis of *Mita*^-/-^ MLFs reconstituted with FLAG-MITA^N153S^ followed by treatment with TAT-WT or TAT-SIP2 (5 μM) for 24 h, and co-localization analysis was conducted. (B) Immunofluorescent staining of Calnexin (an ER marker) (red), GM130 (a Golgi apparatus marker) (red), and FLAG (green) and confocal microscopy analysis of *Mita*^-/-^ MLFs reconstituted with FLAG-MITA followed by transfection with cGAMP (1 μg/mL) for 4 h in the presence of TAT-WT or TAT-SIP2 (5 μM), and co-localization analysis was conducted. (C) Immunofluorescent staining of Calnexin (an ER marker) (red), GM130 (a Golgi apparatus marker) (red), and FLAG (green) and confocal microscopy analysis of *Mita*^-/-^ MLFs reconstituted with FLAG-MITA^R283S^ followed by treatment with TAT-SIP2 (5 μM) for 24 h, and co-localization analysis was conducted. (D) RT-qPCR analysis of *Ifnb*, *Cxcl10*, *Isg56*, *Il6* mRNA levels of MITA^WT/WT^ or MITA^N153S/WT^ MLFs treated with TAT-WT or TAT-SIP2 and followed by treated with HSV-1 for 0-4h. (E) RT-qPCR analysis of *Tnf*, *Ccl2*, *Ccl3* and *Ccl4* mRNA levels in the livers and brains of MITA^N153S/WT^ chimeric mice that were intraperitoneally injected with TAT-WT (n=6) or TAT-SIP2 (n=6) every other day for two weeks starting from the 5th week after bone marrow cell transfer. (F-H) Flow cytometric analysis of the thymocytes (F), bone marrow cells (G) and splenocytes (H) from MITA^N153S/WT^ chimeric mice treated with TAT-WT (n=6) or TAT-SIP2 (n=6) as in (E). \**P*<0.05, \*\**P*<0.01, \*\*\**P*<0.001, \*\*\*\**P*<0.0001; ns, not significant (two-tailed student’s *t*-test). Scale bars represent 10 μm (white) or 1 μm (cyan) in (A-B). The graphs show mean ± SD. The data are representative of three (A-C) or two (D-G) independent experiments.

## References

Abid, Q., Best Rocha, A., Larsen, C.P., Schulert, G., Marsh, R., Yasin, S., Patty-Resk, C., Valentini, R.P., Adams, M., and Baracco, R. (2020). APOL1-Associated Collapsing Focal Segmental Glomerulosclerosis in a Patient With Stimulator of Interferon Genes (STING)-Associated Vasculopathy With Onset in Infancy (SAVI). Am J Kidney Dis 75, 287–290.

Afonine, P.V., Poon, B.K., Read, R.J., Sobolev, O.V., Terwilliger, T.C., Urzhumtsev, A., and Adams, P.D. (2018). Real-space refinement in PHENIX for cryo-EM and crystallography. Acta Crystallogr D Struct Biol 74, 531–544.

Al-Salihi, M.A., and Lang, P.A. (2020). iRhom2: An Emerging Adaptor Regulating Immunity and Disease. Int J Mol Sci 21, 6570.

Balci, S., Ekinci, R.M.K., de Jesus, A.A., Goldbach-Mansky, R., and Yilmaz, M. (2020). Baricitinib experience on STING-associated vasculopathy with onset in infancy: A representative case from Turkey. Clin Immunol 212, 108273.

Cetin Gedik, K., Lamot, L., Romano, M., Demirkaya, E., Piskin, D., Torreggiani, S., Adang, L.A., Armangue, T., Barchus, K., Cordova, D.R., Crow, Y.J., Dale, R.C., Durrant, K.L., Eleftheriou, D., Fazzi, E.M., Gattorno, M., Gavazzi, F., Hanson, E.P., Lee-Kirsch, M.A., Montealegre Sanchez, G.A., Neven, B., Orcesi, S., Ozen, S., Poli, M.C., Schumacher, E., Tonduti, D., Uss, K., Aletaha, D., Feldman, B.M., Vanderver, A., Brogan, P.A., and Goldbach-Mansky, R. (2022). The 2021 European Alliance of Associations for Rheumatology/American College of Rheumatology points to consider for diagnosis and management of autoinflammatory type I interferonopathies: CANDLE/PRAAS, SAVI and AGS. Ann Rheum Dis 81, 601–613.

Chu, L., Qian, L., Chen, Y., Duan, S., Ding, M., Sun, W., Meng, W., Zhu, J., Wang, Q., Hao, H., Wang, C., and Cui, S. (2024). HERC5-catalyzed ISGylation potentiates cGAS-mediated innate immunity. Cell Rep 43, 113870.

Dai, Y., Liu, X., Zhao, Z., He, J., and Yin, Q. (2020). Stimulator of Interferon Genes-Associated Vasculopathy With Onset in Infancy: A Systematic Review of Case Reports. Front Pediatr 8, 577918.

David, C., and Frémond, M.-L. (2022). Lung Inflammation in STING-Associated Vasculopathy with Onset in Infancy (SAVI). Cells 11, 318.

Decout, A., Katz, J.D., Venkatraman, S., and Ablasser, A. (2021). The cGAS–STING pathway as a therapeutic target in inflammatory diseases. Nat Rev Immunol 21, 548–569.

Dobbs, N., Burnaevskiy, N., Chen, D., Gonugunta, V.K., Alto, N.M., and Yan, N. (2015). STING Activation by Translocation from the ER Is Associated with Infection and Autoinflammatory Disease. Cell Host Microbe 18, 157–168.

Emsley, P., Lohkamp, B., Scott, W.G., and Cowtan, K. (2010). Features and development of Coot. Acta Crystallogr D Biol Crystallogr 66, 486–501.

Ergun, S.L., Fernandez, D., Weiss, T.M., and Li, L. (2019). STING Polymer Structure Reveals Mechanisms for Activation, Hyperactivation, and Inhibition. Cell 178, 290–301.e10.

Fang, R., Jiang, Q., Guan, Y., Gao, P., Zhang, R., Zhao, Z., and Jiang, Z. (2021). Golgi apparatus-synthesized sulfated glycosaminoglycans mediate polymerization and activation of the cGAMP sensor STING. Immunity 54, 962–975.e8.

Frémond, M.-L., Hadchouel, A., Berteloot, L., Melki, I., Bresson, V., Barnabei, L., Jeremiah, N., Belot, A., Bondet, V., Brocq, O., Chan, D., Dagher, R., Dubus, J.-C., Duffy, D., Feuillet-Soummer, S., Fusaro, M., Gattorno, M., Insalaco, A., Jeziorski, E., Kitabayashi, N., Lopez-Corbeto, M., Mazingue, F., Morren, M.-A., Rice, G.I., Rivière, J.G., Seabra, L., Sirvente, J., Soler-Palacin, P., Stremler-Le Bel, N., Thouvenin, G., Thumerelle, C., Van Aerde, E., Volpi, S., Willcocks, S., Wouters, C., Breton, S., Molina, T., Bader-Meunier, B., Moshous, D., Fischer, A., Blanche, S., Rieux-Laucat, F., Crow, Y.J., and Neven, B. (2021). Overview of STING-Associated Vasculopathy with Onset in Infancy (SAVI) Among 21 Patients. J Allergy Clin Immunol Pract 9, 803–818.e11.

Frémond, M.-L., Rodero, M.P., Jeremiah, N., Belot, A., Jeziorski, E., Duffy, D., Bessis, D., Cros, G., Rice, G.I., Charbit, B., Hulin, A., Khoudour, N., Caballero, C.M., Bodemer, C., Fabre, M., Berteloot, L., Le Bourgeois, M., Reix, P., Walzer, T., Moshous, D., Blanche, S., Fischer, A., Bader-Meunier, B., Rieux-Laucat, F., Crow, Y.J., and Neven, B. (2016). Efficacy of the Janus kinase 1/2 inhibitor ruxolitinib in the treatment of vasculopathy associated with TMEM173-activating mutations in 3 children. J Allergy Clin Immunol 138, 1752–1755.

Gao, M., Qi, Y., and Zhang, J. (2025). IRF1 amplifies HSV-1-triggered antiviral innate immunity in a feed-forward manner. Cell Insight 4, 100255.

Gao, P., Ascano, M., Zillinger, T., Wang, W., Dai, P., Serganov, A.A., Gaffney, B.L., Shuman, S., Jones, R.A., Deng, L., Hartmann, G., Barchet, W., Tuschl, T., and Patel, D.J. (2013). Structure-function analysis of STING activation by c[G(2’,5’)pA(3’,5’)p] and targeting by antiviral DMXAA. Cell 154, 748–762.

Guo, Y.-Y., Gao, Y., Hu, Y.-R., Zhao, Y., Jiang, D., Wang, Y., Zhang, Y., Gan, H., Xie, C., Liu, Z., Zhong, B., Zhang, Z.-D., and Yao, J. (2022). The Transient Receptor Potential Vanilloid 2 (TRPV2) Channel Facilitates Virus Infection Through the Ca2+ -LRMDA Axis in Myeloid. Cells Adv Sci (Weinh) 9, e2202857.

Hussain, B., Xie, Y., Jabeen, U., Lu, D., Yang, B., Wu, C., and Shang, G. (2022). Activation of STING Based on Its Structural Features. Front Immunol 13, 808607.

Ishikawa, H., and Barber, G.N. (2008). STING is an endoplasmic reticulum adaptor that facilitates innate immune signalling. Nature 455, 674–678.

Ishikawa, H., Ma, Z., and Barber, G.N. (2009). STING regulates intracellular DNA-mediated, type I interferon-dependent innate immunity. Nature 461, 788–792.

Jeremiah, N., Neven, B., Gentili, M., Callebaut, I., Maschalidi, S., Stolzenberg, M.-C., Goudin, N., Frémond, M.-L., Nitschke, P., Molina, T.J., Blanche, S., Picard, C., Rice, G.I., Crow, Y.J., Manel, N., Fischer, A., Bader-Meunier, B., and Rieux-Laucat, F. (2014). Inherited STING-activating mutation underlies a familial inflammatory syndrome with lupus-like manifestations. J Clin Invest 124, 5516–5520.

Jin, L., Waterman, P.M., Jonscher, K.R., Short, C.M., Reisdorph, N.A., and Cambier, J.C. (2008). MPYS, a novel membrane tetraspanner, is associated with major histocompatibility complex class II and mediates transduction of apoptotic signals. Mol Cell Biol 28, 5014–5026.

Kanazawa, N., Ishii, T., Takita, Y., Nishikawa, A., and Nishikomori, R. (2023). Efficacy and safety of baricitinib in Japanese patients with autoinflammatory type I interferonopathies (NNS/CANDLE, SAVI, And AGS). Pediatr Rheumatol Online J 21, 38.

Keskitalo, S., Haapaniemi, E., Einarsdottir, E., Rajamäki, K., Heikkilä, H., Ilander, M., Pöyhönen, M., Morgunova, E., Hokynar, K., Lagström, S., Kivirikko, S., Mustjoki, S., Eklund, K., Saarela, J., Kere, J., Seppänen, M.R.J., Ranki, A., Hannula-Jouppi, K., and Varjosalo, M. (2019). Novel TMEM173 Mutation and the Role of Disease Modifying Alleles. Front Immunol 10, 2770.

Konno, H., Chinn, I.K., Hong, D., Orange, J.S., Lupski, J.R., Mendoza, A., Pedroza, L.A., and Barber, G.N. (2018). Pro-inflammation Associated with a Gain-of-Function Mutation (R284S) in the Innate Immune Sensor STING. Cell Rep 23, 1112–1123.

Kucukelbir, A., Sigworth, F.J., and Tagare, H.D. (2014). Quantifying the local resolution of cryo-EM density maps. Nat Methods 11, 63–65.

Landman, S.L., Ressing, M.E., and van der Veen, A.G. (2020). Balancing STING in antimicrobial defense and autoinflammation. Cytokine Growth Factor Rev 55, 1–14.

Li, Q., Lin, L., Tong, Y., Liu, Y., Mou, J., Wang, X., Wang, X., Gong, Y., Zhao, Y., Liu, Y., Zhong, B., Dai, L., Wei, Y.-Q., Zhang, H., and Hu, H. (2018a). TRIM29 negatively controls antiviral immune response through targeting STING for degradation. Cell Discov 4, 13.

Li, S., Hong, Z., Wang, Z., Li, F., Mei, J., Huang, L., Lou, X., Zhao, S., Song, L., Chen, W., Wang, Q., Liu, H., Cai, Y., Yu, H., Xu, H., Zeng, G., Wang, Q., Zhu, J., Liu, X., Tan, N., and Wang, C. (2018b). The Cyclopeptide Astin C Specifically Inhibits the Innate Immune CDN Sensor STING. Cell Rep 25, 3405–3421.e7.

Li, W., Wang, W., Wang, W., Zhong, L., Gou, L., Wang, C., Ma, J., Quan, M., Jian, S., Tang, X., Zhang, Y., Wang, L., Ma, M., and Song, H. (2022). Janus Kinase Inhibitors in the Treatment of Type I Interferonopathies: A Case Series From a Single Center in China. Front Immunol 13, 825367.

Li, Y., Chen, R., Zhou, Q., Xu, Z., Li, C., Wang, S., Mao, A., Zhang, X., He, W., and Shu, H.-B. (2012). LSm14A is a processing body-associated sensor of viral nucleic acids that initiates cellular antiviral response in the early phase of viral infection. Proc Natl Acad Sci U S A 109, 11770–11775.

Lin, B., Torreggiani, S., Kahle, D., Rumsey, D.G., Wright, B.L., Montes-Cano, M.A., Silveira, L.F., Alehashemi, S., Mitchell, J., Aue, A.G., Ji, Z., Jin, T., de Jesus, A.A., and Goldbach-Mansky, R. (2021). Case Report: Novel SAVI-Causing Variants in STING1 Expand the Clinical Disease Spectrum and Suggest a Refined Model of STING Activation. Front Immunol 12, 636225.

Lin, C., Kuffour, E.O., Fuchs, N.V., Gertzen, C.G.W., Kaiser, J., Hirschenberger, M., Tang, X., Xu, H.C., Michel, O., Tao, R., Haase, A., Martin, U., Kurz, T., Drexler, I., Görg, B., Lang, P.A., Luedde, T., Sparrer, K.M.J., Gohlke, H., König, R., and Münk, C. (2023). Regulation of STING activity in DNA sensing by ISG15 modification. Cell Rep 42, 113277.

Liu, S., Cai, X., Wu, J., Cong, Q., Chen, X., Li, T., Du, F., Ren, J., Wu, Y.-T., Grishin, N.V., and Chen, Z.J. (2015). Phosphorylation of innate immune adaptor proteins MAVS, STING, and TRIF induces IRF3 activation. Science 347, aaa2630.

Liu, S., Yang, B., Hou, Y., Cui, K., Yang, X., Li, X., Chen, L., Liu, S., Zhang, Z., Jia, Y., Xie, Y., Xue, Y., Li, X., Yan, B., Wu, C., Deng, W., Qi, J., Lu, D., Gao, G.F., Wang, P., and Shang, G. (2023). The mechanism of STING autoinhibition and activation. Molecular Cell 83, 1502–1518.e10.

Liu, Y., Jesus, A.A., Marrero, B., Yang, D., Ramsey, S.E., Sanchez, G.A.M., Tenbrock, K., Wittkowski, H., Jones, O.Y., Kuehn, H.S., Lee, C.-C.R., DiMattia, M.A., Cowen, E.W., Gonzalez, B., Palmer, I., DiGiovanna, J.J., Biancotto, A., Kim, H., Tsai, W.L., Trier, A.M., Huang, Y., Stone, D.L., Hill, S., Kim, H.J., St Hilaire, C., Gurprasad, S., Plass, N., Chapelle, D., Horkayne-Szakaly, I., Foell, D., Barysenka, A., Candotti, F., Holland, S.M., Hughes, J.D., Mehmet, H., Issekutz, A.C., Raffeld, M., McElwee, J., Fontana, J.R., Minniti, C.P., Moir, S., Kastner, D.L., Gadina, M., Steven, A.C., Wingfield, P.T., Brooks, S.R., Rosenzweig, S.D., Fleisher, T.A., Deng, Z., Boehm, M., Paller, A.S., and Goldbach-Mansky, R. (2014). Activated STING in a vascular and pulmonary syndrome. N Engl J Med 371, 507–518.

Lu, D., Shang, G., Zhang, H., Yu, Q., Cong, X., Yuan, J., He, F., Zhu, C., Zhao, Y., Yin, K., Chen, Y., Hu, J., Zhang, X., Yuan, Z., Xu, S., Hu, W., Cang, H., and Gu, L. (2014). Structural insights into the T6SS effector protein Tse3 and the Tse3-Tsi3 complex from Pseudomonas aeruginosa reveal a calcium-dependent membrane-binding mechanism. Mol Microbiol 92, 1092–1112.

Luksch, H., Stinson, W.A., Platt, D.J., Qian, W., Kalugotla, G., Miner, C.A., Bennion, B.G., Gerbaulet, A., Rösen-Wolff, A., and Miner, J.J. (2019). STING-associated lung disease in mice relies on T cells but not type I interferon. J Allergy Clin Immunol 144, 254–266.e8.

Luo, W.-W., Li, S., Li, C., Lian, H., Yang, Q., Zhong, B., and Shu, H.-B. (2016). iRhom2 is essential for innate immunity to DNA viruses by mediating trafficking and stability of the adaptor STING. Nat Immunol 17, 1057–1066.

Luo, W.-W., Li, S., Li, C., Zheng, Z.-Q., Cao, P., Tong, Z., Lian, H., Wang, S.-Y., Shu, H.-B., and Wang, Y.-Y. (2017). iRhom2 is essential for innate immunity to RNA virus by antagonizing ER- and mitochondria-associated degradation of VISA. PLoS Pathog 13, e1006693.

Melki, I., Rose, Y., Uggenti, C., Van Eyck, L., Frémond, M.-L., Kitabayashi, N., Rice, G.I., Jenkinson, E.M., Boulai, A., Jeremiah, N., Gattorno, M., Volpi, S., Sacco, O., Terheggen-Lagro, S.W.J., Tiddens, H.A.W.M., Meyts, I., Morren, M.-A., De Haes, P., Wouters, C., Legius, E., Corveleyn, A., Rieux-Laucat, F., Bodemer, C., Callebaut, I., Rodero, M.P., and Crow, Y.J. (2017). Disease-associated mutations identify a novel region in human STING necessary for the control of type I interferon signaling. J Allergy Clin Immunol 140, 543–552.e5.

Meng, E.C., Goddard, T.D., Pettersen, E.F., Couch, G.S., Pearson, Z.J., Morris, J.H., and Ferrin, T.E. (2023). UCSF ChimeraX: Tools for structure building and analysis. Protein Sci 32, e4792.

Miner, J.J., and Fitzgerald, K.A. (2023). A path towards personalized medicine for autoinflammatory and related diseases. Nat Rev Rheumatol 19, 182–189.

Mitrovic, S., Ben-Tekaya, H., Koegler, E., Gruenberg, J., and Hauri, H.-P. (2008). The cargo receptors Surf4, endoplasmic reticulum-Golgi intermediate compartment (ERGIC)-53, and p25 are required to maintain the architecture of ERGIC and Golgi. Mol Biol Cell 19, 1976–1990.

MotionCor2: anisotropic correction of beam-induced motion for improved cryo-electron microscopy - PubMed (n.d.). URL https://pubmed.ncbi.nlm.nih.gov/28250466/ (accessed 10 July 2025).

Motwani, M., Pawaria, S., Bernier, J., Moses, S., Henry, K., Fang, T., Burkly, L., Marshak-Rothstein, A., and Fitzgerald, K.A. (2019). Hierarchy of clinical manifestations in SAVI N153S and V154M mouse models Proc. Natl. Acad. Sci. U.S.A. 116, 7941–7950.

Mukai, K., Ogawa, E., Uematsu, R., Kuchitsu, Y., Kiku, F., Uemura, T., Waguri, S., Suzuki, T., Dohmae, N., Arai, H., Shum, A.K., and Taguchi, T. (2021). Homeostatic regulation of STING by retrograde membrane traffic to the ER. Nat Commun 12, 61.

Munoz, J., Rodière, M., Jeremiah, N., Rieux-Laucat, F., Oojageer, A., Rice, G.I., Rozenberg, F., Crow, Y.J., and Bessis, D. (2015). Stimulator of Interferon Genes-Associated Vasculopathy With Onset in Infancy: A Mimic of Childhood Granulomatosis With Polyangiitis. JAMA Dermatol 151, 872–877.

Pettersen, E.F., Goddard, T.D., Huang, C.C., Couch, G.S., Greenblatt, D.M., Meng, E.C., and Ferrin, T.E. (2004). UCSF Chimera--a visualization system for exploratory research and analysis. J Comput Chem 25, 1605–1612.

Punjani, A., Rubinstein, J.L., Fleet, D.J., and Brubaker, M.A. (2017). cryoSPARC: algorithms for rapid unsupervised cryo-EM structure determination. Nat Methods 14, 290–296.

Pymol: an open-source molecular graphics tool – ScienceOpen (n.d.). URL https://www.scienceopen.com/document?vid=4362f9a2-0b29-433f-aa65-51db01f4962f (accessed 10 July 2025).

Qin, Y., Wang, M., Meng, X., Wang, M., Jiang, H., Gao, Y., Li, J., Zhao, C., Han, C., Zhao, W., and Zheng, X. (2024). ISGylation by HERCs facilitates STING activation. Cell Rep 43, 114135.

Qin, Y., Zhou, M.-T., Hu, M.-M., Hu, Y.-H., Zhang, J., Guo, L., Zhong, B., and Shu, H.-B. (2014). RNF26 temporally regulates virus-triggered type I interferon induction by two distinct mechanisms. PLoS Pathog 10, e1004358.

Qing, X., Chinenov, Y., Redecha, P., Madaio, M., Roelofs, J.J., Farber, G., Issuree, P.D., Donlin, L., Mcllwain, D.R., Mak, T.W., Blobel, C.P., and Salmon, J.E. (2018). iRhom2 promotes lupus nephritis through TNF-α and EGFR signaling. J Clin Invest 128, 1397–1412.

Rohou, A., and Grigorieff, N. (2015). CTFFIND4: Fast and accurate defocus estimation from electron micrographs. J Struct Biol 192, 216–221.

Saldanha, R.G., Balka, K.R., Davidson, S., Wainstein, B.K., Wong, M., Macintosh, R., Loo, C.K.C., Weber, M.A., Kamath, V., Circa, Aadry, Moghaddas, F., De Nardo, D., Gray, P.E., and Masters, S.L. (2018). A Mutation Outside the Dimerization Domain Causing Atypical STING-Associated Vasculopathy With Onset in Infancy. Front Immunol 9, 1535.

Sanchez, G.A.M., Reinhardt, A., Ramsey, S., Wittkowski, H., Hashkes, P.J., Berkun, Y., Schalm, S., Murias, S., Dare, J.A., Brown, D., Stone, D.L., Gao, L., Klausmeier, T., Foell, D., de Jesus, A.A., Chapelle, D.C., Kim, H., Dill, S., Colbert, R.A., Failla, L., Kost, B., O’Brien, M., Reynolds, J.C., Folio, L.R., Calvo, K.R., Paul, S.M., Weir, N., Brofferio, A., Soldatos, A., Biancotto, A., Cowen, E.W., Digiovanna, J.J., Gadina, M., Lipton, A.J., Hadigan, C., Holland, S.M., Fontana, J., Alawad, A.S., Brown, R.J., Rother, K.I., Heller, T., Brooks, K.M., Kumar, P., Brooks, S.R., Waldman, M., Singh, H.K., Nickeleit, V., Silk, M., Prakash, A., Janes, J.M., Ozen, S., Wakim, P.G., Brogan, P.A., Macias, W.L., and Goldbach-Mansky, R. (2018). JAK1/2 inhibition with baricitinib in the treatment of autoinflammatory interferonopathies. J Clin Invest 128, 3041–3052.

Seo, G.J., Kim, C., Shin, W.-J., Sklan, E.H., Eoh, H., and Jung, J.U. (2018). TRIM56-mediated monoubiquitination of cGAS for cytosolic DNA sensing. Nat Commun 9, 613.

Seo, J., Kang, J.-A., Suh, D.I., Park, E.-B., Lee, C.-R., Choi, S.A., Kim, S.Y., Kim, Y., Park, S.-H., Ye, M., Kwon, S.-H., Park, J.D., Lim, B.C., Lee, D.H., Kang, S.-J., Choi, M., Park, S.-G., and Chae, J.-H. (2017). Tofacitinib relieves symptoms of stimulator of interferon genes (STING)-associated vasculopathy with onset in infancy caused by 2 de novo variants in TMEM173. J Allergy Clin Immunol 139, 1396–1399.e12.

Shang, G., Zhang, C., Chen, Z.J., Bai, X.-C., and Zhang, X. (2019). Cryo-EM structures of STING reveal its mechanism of activation by cyclic GMP-AMP. Nature 567, 389–393.

Shen, Y., Gu, H.-M., Qin, S., and Zhang, D.-W. (2023). Surf4, cargo trafficking, lipid metabolism, and therapeutic implications. J Mol Cell Biol 14, mjac063.

Shi, H.-X., Yang, K., Liu, X., Liu, X.-Y., Wei, B., Shan, Y.-F., Zhu, L.-H., and Wang, C. (2010). Positive regulation of interferon regulatory factor 3 activation by Herc5 via ISG15 modification. Mol Cell Biol 30, 2424–2436.

Srikanth, S., Woo, J.S., Wu, B., El-Sherbiny, Y.M., Leung, J., Chupradit, K., Rice, L., Seo, G.J., Calmettes, G., Ramakrishna, C., Cantin, E., An, D.S., Sun, R., Wu, T.-T., Jung, J.U., Savic, S., and Gwack, Y. (2019). The Ca2+ sensor STIM1 regulates the type I interferon response by retaining the signaling adaptor STING at the endoplasmic reticulum. Nat Immunol 20, 152–162.

Sun, H., Zhang, Q., Jing, Y.-Y., Zhang, M., Wang, H.-Y., Cai, Z., Liuyu, T., Zhang, Z.-D., Xiong, T.-C., Wu, Y., Zhu, Q.-Y., Yao, J., Shu, H.-B., Lin, D., and Zhong, B. (2017). USP13 negatively regulates antiviral responses by deubiquitinating STING. Nat Commun 8, 15534.

Sun, L., Wu, J., Du, F., Chen, X., and Chen, Z.J. (2013). Cyclic GMP-AMP synthase is a cytosolic DNA sensor that activates the type I interferon pathway. Science 339, 786–791.

Sun, M.-S., Zhang, J., Jiang, L.-Q., Pan, Y.-X., Tan, J.-Y., Yu, F., Guo, L., Yin, L., Shen, C., Shu, H.-B., and Liu, Y. (2018). TMED2 Potentiates Cellular IFN Responses to DNA Viruses by Reinforcing MITA Dimerization and Facilitating Its Trafficking. Cell Reports 25, 3086–3098.e3.

Sun, W., Li, Y., Chen, L., Chen, H., You, F., Zhou, X., Zhou, Y., Zhai, Z., Chen, D., and Jiang, Z. (2009). ERIS, an endoplasmic reticulum IFN stimulator, activates innate immune signaling through dimerization. Proc Natl Acad Sci U S A 106, 8653–8658.

Tanaka, Y., and Chen, Z.J. (2012). STING specifies IRF3 phosphorylation by TBK1 in the cytosolic DNA signaling pathway. Sci Signal 5, ra20.

Tang, X., Xu, H., Zhou, C., Peng, Y., Liu, H., Liu, J., Li, H., Yang, H., and Zhao, S. (2020). STING-Associated Vasculopathy with Onset in Infancy in Three Children with New Clinical Aspect and Unsatisfactory Therapeutic Responses to Tofacitinib. J Clin Immunol 40, 114–122.

Tokgun, P.E., Karagenc, N., Karasu, U., Tokgun, O., Turel, S., Demiray, A., Akca, H., and Yüksel, S. (2023). Treatment of STING-associated vasculopathy with onset in infancy in patients carrying a novel mutation in the TMEM173 gene with the JAK3-inhibitor tofacitinib. Arch Rheumatol 38, 461–467.

Tsuchida, T., Zou, J., Saitoh, T., Kumar, H., Abe, T., Matsuura, Y., Kawai, T., and Akira, S. (2010). The Ubiquitin Ligase TRIM56 Regulates Innate Immune Responses to Intracellular Double-Stranded DNA. Immunity 33, 765–776.

Valeri, E., Breggion, S., Barzaghi, F., Abou Alezz, M., Crivicich, G., Pagani, I., Forneris, F., Sartirana, C., Costantini, M., Costi, S., Marino, A., Chiarotto, E., Colavito, D., Cimaz, R., Merelli, I., Vicenzi, E., Aiuti, A., and Kajaste-Rudnitski, A. (2024). A novel STING variant triggers endothelial toxicity and SAVI disease. J Exp Med 221, e20232167.

Volpi, S., Insalaco, A., Caorsi, R., Santori, E., Messia, V., Sacco, O., Terheggen-Lagro, S., Cardinale, F., Scarselli, A., Pastorino, C., Moneta, G., Cangemi, G., Passarelli, C., Ricci, M., Girosi, D., Derchi, M., Bocca, P., Diociaiuti, A., El Hachem, M., Cancrini, C., Tomà, P., Granata, C., Ravelli, A., Candotti, F., Picco, P., DeBenedetti, F., and Gattorno, M. (2019). Efficacy and Adverse Events During Janus Kinase Inhibitor Treatment of SAVI Syndrome. J Clin Immunol 39, 476–485.

Wan, R., Fänder, J., Zakaraia, I., Lee-Kirsch, M.A., Wolf, C., Lucas, N., Olfe, L.I., Hendrich, C., Jonigk, D., Holzinger, D., Steindor, M., Schmidt, G., Davenport, C., Klemann, C., Schwerk, N., Griese, M., Schlegelberger, B., Stehling, F., Happle, C., Auber, B., Steinemann, D., Wetzke, M., and von Hardenberg, S. (2022). Phenotypic spectrum in recessive STING-associated vasculopathy with onset in infancy: Four novel cases and analysis of previously reported cases. Front Immunol 13, 1029423.

Wang, Y., Wang, F., and Zhang, X. (2021). STING-associated vasculopathy with onset in infancy: a familial case series report and literature review. Ann Transl Med 9, 176–176.

Warner, J.D., Irizarry-Caro, R.A., Bennion, B.G., Ai, T.L., Smith, A.M., Miner, C.A., Sakai, T., Gonugunta, V.K., Wu, J., Platt, D.J., Yan, N., and Miner, J.J. (2017). STING-associated vasculopathy develops independently of IRF3 in mice. Journal of Experimental Medicine 214, 3279–3292.

Williams, C.J., Headd, J.J., Moriarty, N.W., Prisant, M.G., Videau, L.L., Deis, L.N., Verma, V., Keedy, D.A., Hintze, B.J., Chen, V.B., Jain, S., Lewis, S.M., Arendall, W.B., Snoeyink, J., Adams, P.D., Lovell, S.C., Richardson, J.S., and Richardson, D.C. (2018). MolProbity: More and better reference data for improved all-atom structure validation. Protein Sci 27, 293–315.

Wu, J., Sun, L., Chen, X., Du, F., Shi, H., Chen, C., and Chen, Z.J. (2013). Cyclic GMP-AMP is an endogenous second messenger in innate immune signaling by cytosolic. DNA Science 339, 826–830.

Xing, J., Zhang, A., Zhang, H., Wang, J., Li, X.C., Zeng, M.-S., and Zhang, Z. (2017). TRIM29 promotes DNA virus infections by inhibiting innate immune response. Nat Commun 8, 945.

Xu, Q., Chen, C., Liu, B., Lin, Y., Zheng, P., Zhou, D., Xie, Y., Lin, Y., Guo, C., Liu, J., and Li, L. (2020). Association of iRhom1 and iRhom2 expression with prognosis in patients with cervical cancer and possible signaling pathways. Oncol Rep 43, 41–54.

Yang, M., Chen, X., Hu, X., Li, H., Huang, H., Fang, Y., Jiang, J., Liu, H., Wang, Y., and Qing, G. (2025). The NF-κB-SLC7A11 axis regulates ferroptosis sensitivity in inflammatory macrophages Cell Insight 4, 100257.

Zeng, J.-Q., Ruan, Z.-L., Zhang, Q., Yi, X.-M., Chen, Y.-D., Hu, M.-M., and Li, S. (2026). HTATSF1 regulates innate antiviral immune response by orchestrating TRAF3-IRF3 and TRAF6-NF-κB pathways. Cell Insight 5, 100294.

Zhang, B.-C., Nandakumar, R., Reinert, L.S., Huang, J., Laustsen, A., Gao, Z.-L., Sun, C.-L., Jensen, S.B., Troldborg, A., Assil, S., Berthelsen, M.F., Scavenius, C., Zhang, Y., Windross, S.J., Olagnier, D., Prabakaran, T., Bodda, C., Narita, R., Cai, Y., Zhang, C.-G., Stenmark, H., Doucet, C.M., Noda, T., Guo, Z., Goldbach-Mansky, R., Hartmann, R., Chen, Z.J., Enghild, J.J., Bak, R.O., Thomsen, M.K., and Paludan, S.R. (2020a). STEEP mediates STING ER exit and activation of signaling. Nat Immunol 21, 868–879.

Zhang, C., Shang, G., Gui, X., Zhang, X., Bai, X.-C., and Chen, Z.J. (2019). Structural basis of STING binding with and phosphorylation by TBK1. Nature 567, 394–398.

Zhang, X., Shi, H., Wu, J., Zhang, X., Sun, L., Chen, C., and Chen, Z.J. (2013). Cyclic GMP-AMP containing mixed phosphodiester linkages is an endogenous high-affinity ligand for STING. Mol Cell 51, 226–235.

Zhang, Z.-D., Li, H.-X., Gan, H., Tang, Z., Guo, Y.-Y., Yao, S.-Q., Liuyu, T., Zhong, B., and Lin, D. (2022). RNF115 Inhibits the Post-ER Trafficking of TLRs and TLRs-Mediated Immune Responses by Catalyzing K11-Linked Ubiquitination of RAB1A and RAB13. Adv Sci (Weinh) 9, e2105391.

Zhang, Z.-D., Shi, C.-R., Li, F.-X., Gan, H., Wei, Y., Zhang, Q., Shuai, X., Chen, M., Lin, Y.-L., Xiong, T.-C., Chen, X., Zhong, B., and Lin, D. (2024). Disulfiram ameliorates STING/MITA-dependent inflammation and autoimmunity by targeting RNF115. Cell Mol Immunol.

Zhang, Z.-D., Xiong, T.-C., Yao, S.-Q., Wei, M.-C., Chen, M., Lin, D., and Zhong, B. (2020). RNF115 plays dual roles in innate antiviral responses by catalyzing distinct ubiquitination of MAVS and MITA. Nat Commun 11, 5536.

Zhong, B., and Shu, H.-B. (2022). MITA/STING-mediated antiviral immunity and autoimmunity: the evolution, mechanism, and intervention. Curr Opin Immunol 78, 102248.

Zhong, B., Yang, Y., Li, S., Wang, Y.-Y., Li, Y., Diao, F., Lei, C., He, X., Zhang, L., Tien, P., and Shu, H.-B. (2008). The adaptor protein MITA links virus-sensing receptors to IRF3 transcription factor activation. Immunity 29, 538–550.

Zhong, B., Zhang, L., Lei, C., Li, Y., Mao, A.-P., Yang, Y., Wang, Y.-Y., Zhang, X.-L., and Shu, H.-B. (2009). The ubiquitin ligase RNF5 regulates antiviral responses by mediating degradation of the adaptor protein MITA. Immunity 30, 397–407.

Zivanov, J., Nakane, T., Forsberg, B.O., Kimanius, D., Hagen, W.J., Lindahl, E., and Scheres, S.H. (2018). New tools for automated high-resolution cryo-EM structure determination in RELION-3. Elife 7, e42166.

